# Single-cell trait diversity explains niche and fitness differences in aquatic microbial communities

**DOI:** 10.1101/2024.02.07.579265

**Authors:** Simone Fontana, Désirée A. Schmitz, Michael Daniels, Francesco Danza, Thomas Röösli, Hannah Bruderer, Jean-Claude Walser, Julien Dekaezemacker, Stephane Escrig, Anders Meibom, Francesco Pomati, Frank Schreiber

## Abstract

Two fundamental questions in ecology are how biodiversity is maintained and how it affects ecosystem functioning. Until now, it has been difficult to study the above mechanisms in natural microbial communities, yet they are important drivers of biogeochemical ecosystem functions. Here, we use a new approach to define and measure biodiversity in complex lake microbial communities (Lake Cadagno, Switzerland) based on the cell-to-cell variation in multiple functionally relevant phenotypic traits. We use stable isotope probing coupled to correlative imaging using confocal laser scanning microscopy (CLSM) and nanometer-scale secondary ion mass spectrometry (NanoSIMS) to obtain morphological (size), physiological (pigments) and metabolic (carbon and nitrogen isotope uptake and sulfur content) traits for a large number of individual cells along the environmental gradient found across lake depth. We show that cell-to-cell trait variation is significantly correlated with cell densities as a proxy for ecosystem functioning, whereas genetic diversity measured at the level of 16S and 18S is not. Our single-cell analysis provides evidence for a simultaneous increase in niche partitioning (measured as increased evenness in pigment composition) and decrease in fitness differences (measured as decreased variability in sulfur content) due to light limitation and competition for nutrients in deep layers of the lake. This leads to a negative relationship between niche and fitness differences. Our results suggest that niche and fitness differences in natural microbial communities can be understood at the level of single-cell traits, providing a mechanistic understanding of the relationship between microbial diversity and ecosystem functioning.

## Introduction

Modern and past biodiversity levels on Earth have raised two central questions in ecological research: how did this diversity arise and how is it maintained in natural ecosystems? The principle of competitive exclusion^1–3^ and the closely related concept of ecological niche^4–6^ initially led to the recognition that species have different requirements and use distinct resources, thus explaining their coexistence. However, the number of different types of organisms – especially in microbial communities – greatly exceeds the number of distinct limiting resources (e.g., ‘the paradox of the plankton’^7^), and other mechanisms must also play a role. These mechanisms have broadly been divided into two categories: increasing niche partitioning and decreasing fitness differences^8–10^, which are not mutually exclusive. Indeed, both niche and fitness effects can act simultaneously^11,12^ and be considered interdependent^13^. In addition, evolutionary (genetic) mechanisms also act as sources of diversity in microbial communities^14,15^. For example, rapid evolution has been shown to maintain taxonomic diversity in aquatic communities, even in the absence of niche differentiation^16,17^.

In addition to the drivers of diversity, ecological research has also addressed its consequences, i.e., biodiversity and ecosystem functioning relationships (e.g.^18,19^). The mechanisms that generate diversity are arguably related to those determining ecosystem functioning^20–22^. As microbial communities are fundamental drivers of biogeochemical functions, such as the remineralization of polymers and the nitrogen cycle, it is essential to elucidate these mechanisms to better understand the contribution of microbial communities to ecosystem functioning.

Interestingly, both niche partitioning and fitness differences can be the outcome of variation in functional traits that, by definition, are features measurable at the level of individual organisms^23^. In other words, the outcome(s) of coexistence observed at the community level are in fact the result of interactions occurring among individuals^24^. However, studies of niche and fitness differences at the level of the effective competitive units, i.e., individual organisms are still lacking. Addressing this gap is particularly challenging for microorganisms in the natural environment, as it is difficult to measure their relevant traits at the single-cell level. Most research to date in this field has used low specificity measures of trait diversity, typically including multiple traits that provide limited mechanistic understanding, such as broad classes of functionally unrelated genes from metagenomic datasets^25^. However, the advantage of this approach is that it often provides good predictive power of ecosystem functioning^24^. We capitalize on recent advances in the understanding of microbial function through single-cell trait-based approaches to investigate niche and fitness differences in microbial communities based on single-cell phenotypes.

Modern high-throughput single-cell technologies allow for a more mechanistic and hypothesis-driven investigation of microbial adaptive strategies based on individual-level traits^26^. The integration of such technologies has the potential to reveal niche and fitness differences at the level of the individual organisms and assess whether these traits drive stabilizing and equalizing mechanisms at the population and community level. First, the definition of microbial species (assuming they exist) is challenging and not trivial^27–29^. Second, the extreme phenotypic variability observed even within clonal microbial populations^30^ or closely related sub-taxonomic groups^31^ might jeopardize the promise of modern taxonomic approaches linked to rRNA gene sequencing to elucidate relationships between biodiversity and ecosystem function^32–34^. Taken together, a single-cell perspective will improve our understanding of how niche and fitness differences relate to ecosystem functioning in nature, with possible implications for the monitoring of biodiversity and related functions.

In this study, we aim to investigate niche and fitness differences within a natural microbial community and their relationship with ecosystem functioning based on the characteristics of individual microbial cells. In this context, it is crucial to identify which traits might cause niche partitioning, and which may be used to quantify fitness differences. We use correlative fluorescence microscopy and stable isotope probing coupled to nanometer-scale secondary ion mass spectrometry (NanoSIMS)^35^ to determine photosynthetic pigment composition, size, and metabolic activity for carbon, nitrogen and sulfur simultaneously in thousands of individual microbial cells from different samples. We leverage the idea that environmental gradients explain the diversification of ecological strategies or niches^36^, as well as fitness differences between phenotypes^37^. On the one hand, pigment-related traits (i.e., photosynthesis/physiology) enable niche partitioning, as cells can decrease competition for light by targeting slightly different wavelengths through fine-tuning of the relative expression of different pigments^38^. On the other hand, traits defining the assimilation or cell content of elements involved in fundamental metabolic pathways (C, N, and S) - which often show great intraspecific variation^39–41^ - can be considered as proxies for fitness, as they influence the maintenance, growth and reproduction of organisms^42,43^. Here, we sampled single cells living in structured microbial communities at different depths in the meromictic Lake Cadagno (Switzerland), and considered decreasing levels of trait integration to predict ecosystem functioning with multidimensional trait diversity indices and detect niche and fitness differences determined by the variation in physiological and metabolic traits^24^. Accordingly, we formulate the following expectations:

**1)** Ecosystem functioning (i.e., cell density as a proxy for primary production) is better explained by trait diversity metrics that integrate all available individual-based traits (morphological, physiological, and metabolic) rather than by taxonomic diversity metrics based on rRNA gene sequencing^33,44^. **2)** In deep layers of the lake, competition for light drives increasing niche partitioning in pigment-related traits (i.e., increasing trait evenness^38^), while limitation for essential nutrients mediates decreasing fitness differences in metabolic traits (i.e., trait convergence^45^), as compared to the upper layers of the lake. **3)** These simultaneous shifts in trait distribution lead to a negative relationship between single-cell niche and fitness differences across water depth.

## Results

### Environmental variables structure Lake Cadagno into distinct layers

First, we set out to characterize the environmental variables that shape the microbial communities in Lake Cadagno and that potentially drive trait variability, mediating niche and fitness differences. The vertical structure of physical, chemical and biological parameters of the meromictic Lake Cadagno during one day of our sampling campaign is shown in Fig. 1A (see Extended Data Fig. 1 for the profiles of the other two days). Lake Cadagno is perfectly suited to study niche and fitness differences in microbial communities because its steep gradients of light intensity, oxygen and sulfide concentration (Fig. 1A) create ideal conditions for very distinct microbial communities at different depths^46^. Differences in salt concentration stably stratify the lake into three main layers. The upper layer, called mixolimnion, is dominated by oxygenic photosynthetic organisms. Oxygen is consumed around 11 m water depth, below which light can still penetrate. A chemocline then forms in this largely anoxic water, which is dominated by high cell densities of anoxygenic phototrophic purple and green sulfur bacteria. As electron donors, these bacteria use sulfide and other reduced sulfur compounds that diffuse upward from the lake’s bottom waters (Fig. 1A; see also^47^). Below the chemocline, in the bottom waters, light penetration is no longer measurable, and sulfide is produced by sulfate-reducing bacteria (Fig. 1A).

**Fig. 1.**
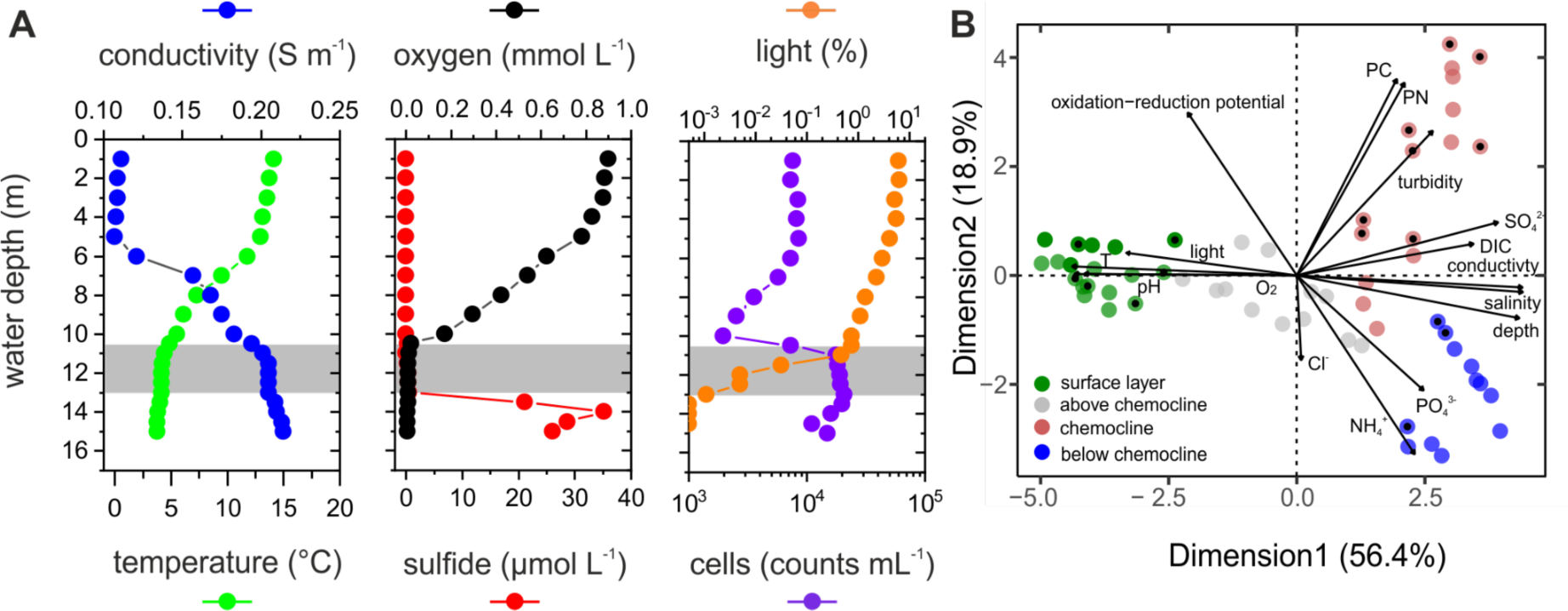
Vertical gradients of physical, chemical and biological parameters in Lake Cadagno separate the ecosystem into three distinct layers. **(A)** Vertical profiles of conductivity (blue), temperature (green), oxygen (O_2_, black), total sulfide (red), light (orange), and cell density (purple) measured on day C (19/08/2014) of the sampling campaign. Profiles from the other sampling days are shown in Extended Data Fig. 1. The gray area represents the chemocline, where sulfide and O_2_ are depleted, the cell density increases, and light gets severely limited. Light values are presented as % photosynthetic active radiation of the light above the water. Light values shown as half circles below the chemocline were below the detection limit of the sensor. **(B)** Principal component analysis (PCA) of individual samples based on environmental factors (abbreviations: T=temperature; Cl^-^=chloride; NH_4_^+^=ammonium; PO_4_^3-^=ortho-phosphate; DIC=dissolved inorganic carbon; SO ^2-^; sulfate; PN=particulate nitrogen; PC=particulate carbon). The PCA plot shows individual samples (circles) taken at different days distributed across environmental factors. Samples belonging to the same sampling layer (color) are grouped more closely in two-dimensional space. Samples for which individual-based trait diversity (*‘Trait Onion Peeling’* TOP, *‘Trait Even Distribution’* TED and *‘Functional Dispersion’* FDis) were calculated for further analysis are marked by a black point. The surface layer is defined hereafter as ‘upper layer’ (as the gray dots are not considered for further analyses), and the layer below chemocline as ‘lower layer’.

Our analysis of the lake stratification based on environmental factors (Fig. 1B) shows that Lake Cadagno is separated into distinct layers and that different environmental factors shape the structure and function of the microbial communities in each layer. Specifically, the upper layer is strongly separated from the chemocline and the lower layer by one axis of the principal component analysis (PCA; dimension 1), which explains 56.4% of the variance. The variables that most strongly contribute to variation in this dimension are depth and light intensity. This indicates that the microbial communities found in these samples are, as expected, stratified based on the availability of light as a resource for photosynthesis. Samples in the chemocline are most strongly separated from the lower layer by other environmental parameters (dimension 2 of the PCA, explaining 18.9% of the variance): high turbidity, oxidation-reduction potential, particulate carbon, and ammonium concentration. Samples in the lower layer correlate most strongly with the concentration of ammonium and ortho-phosphate. These environmental factors thus separate Lake Cadagno into different layers and provide the physio-chemical basis for the structuring of the lake’s microbial communities.

### Single-cell trait diversity explains ecosystem functioning

Next, we tested whether ecosystem functioning can be better explained by integrative trait diversity metrics (including physiological and morphological traits from CLSM and metabolic traits from NanoSIMS; Fig. 2) than taxonomic diversity metrics based on rRNA gene sequencing. To do that, we analyzed trait and taxonomic diversity data across the different depths of the lake and time (days) with linear models to assess their explanatory power for ecosystem functioning, represented here by the density of cells as a proxy for primary production. The analysis of our large dataset across time and depths shows that trait diversity is a significant driver of ecosystem functioning. Specifically, the relative regularity in distances between individual cells (TED, *‘Trait Even Distribution’*) and their spread around the average phenotype (FDis, *‘Functional Dispersion’*) in high-dimensional trait space was significantly related to ecosystem functioning, whereas the total trait space covered (TOP, *‘Trait Onion Peeling’*) did not. Thus, the two best models retained for model averaging explained 56.8% (explanatory variables: TED, FDis) and 71.9% (explanatory variables: TED, FDis, Day of sampling) of the variance in cell density relative to the maximum achieved in a given layer (Extended Data Table S1). Model averaging showed a negative effect of TED on cell density (standardized regression coefficient=-0.195; 95% CI=[-0.339; −0.050]), whereas FDis had a positive effect of similar strength (standardized regression coefficient=0.153; 95% CI=[0.028; 0.279]) (Fig. 3). Contrary to this, taxonomic diversity metrics based on rRNA gene sequencing did not show any significant correlation with cell density. Indeed, the best model according to AICc (Akaike information criterion corrected for small sample sizes) was the null model (including the intercept only). The only other model retained for model averaging also included the Simpson index based on 18S sequencing and only explained 5.7% of the variance in cell density relative to the maximum achieved in a given layer (Extended Data Table S1). As a consequence, the effect size of the Simpson index based on 18S sequencing is not significantly different from 0 (standardized regression coefficient=0.026; 95% CI=[-0.079; 0.130]) (Fig. 3). Taken together, these data show that trait diversity explains ecosystem functioning in Lake Cadagno while taxonomic diversity does not.

**Fig. 2.**
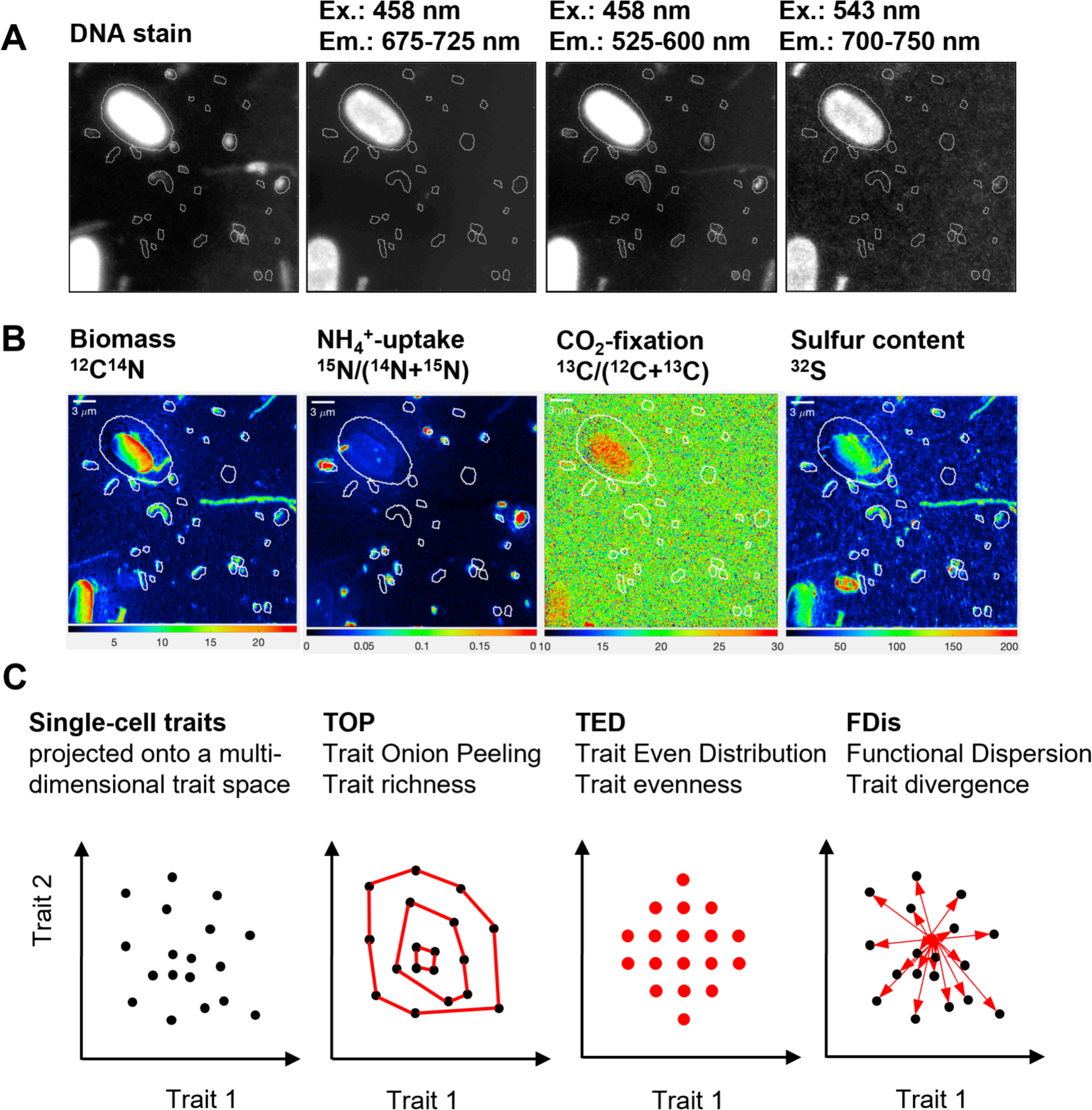
Correlative microscopy to determine single-cell trait-based diversity indices. Panels **(A)** and **(B)** show example images for the same region of interest (ROI) measured with confocal laser scanning microscopy and NanoSIMS, respectively. The lines around cells show the cell segmentation that was based on DNA stain and the ^12^C^14^N NanoSIMS signal. Cells that are at the edge of each ROI were not included in the analysis. The excitation (Ex.) and emission (Em.) wavelengths are shown above each microscopy image. The intensity scale for isotope counts (^12^C^14^N and ^32^S) and isotope fractions (^15^N/[^14^N+^15^N] and ^13^C/[^12^C+^13^C]) are shown below each NanoSIMS image. Scale bars are only shown in the NanoSIMS images, but they are also valid for the microscopy images since they show the same ROI. **(C)** Scheme summarizing how the measured single-cell traits are used to calculate the three components of trait diversity: richness (TOP index), evenness (TED index) and divergence (FDis index). Note that the scheme shows the indices on a 2-dimensional trait axis to better understand the principle. In this study, the trait axes form a multi-dimensional trait space with seven dimensions represented by fluorescence data (three channels) and cell size from CLSM, and NanoSIMS data (C and N assimilation, as well as S cell content).

**Fig. 3:**
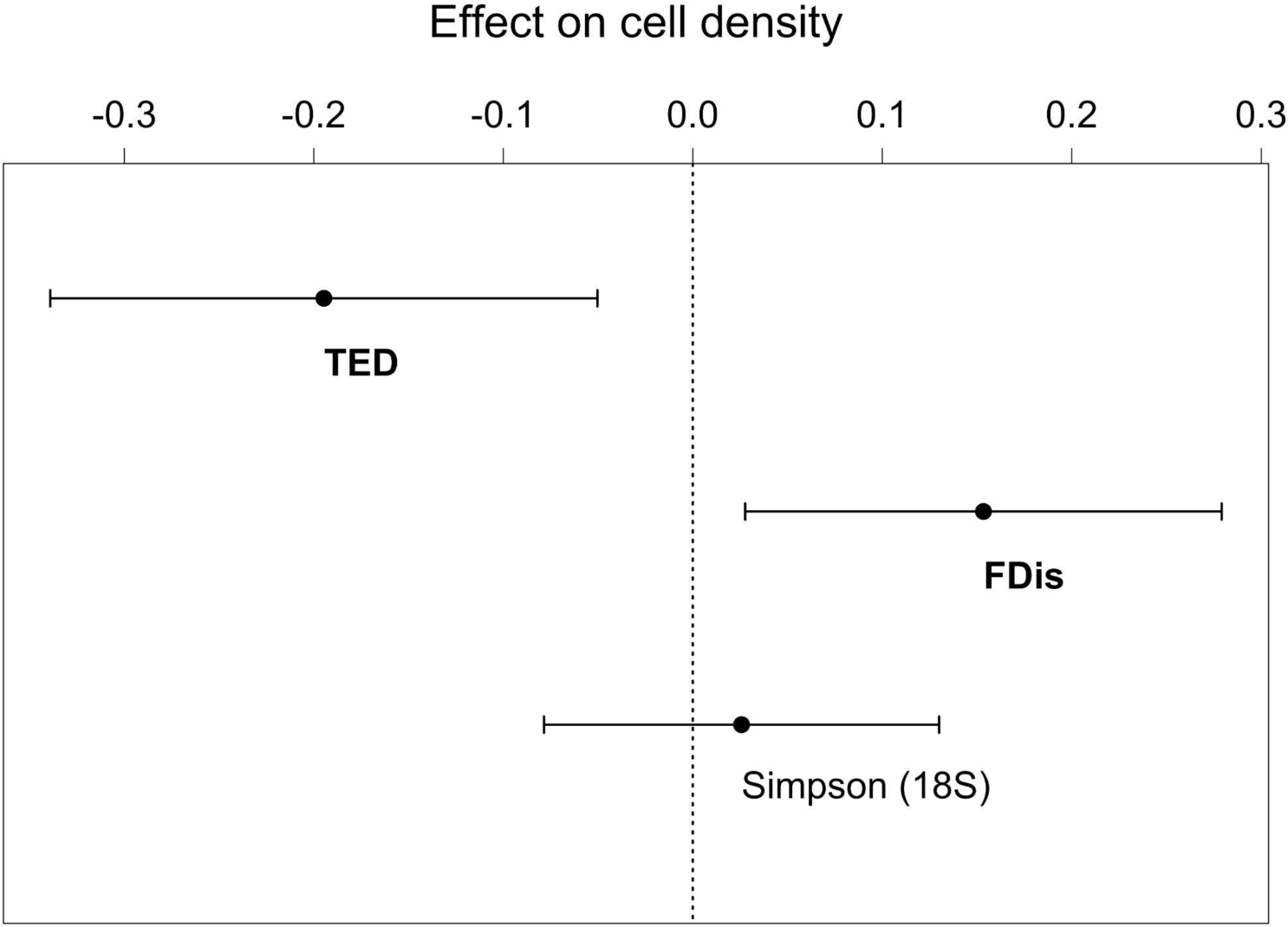
Single-cell-based trait diversity explains ecosystem functioning in Lake Cadagno, whereas taxonomic diversity based on rRNA gene sequencing does not. Effect size of trait diversity metrics (TOP index of trait richness, TED index of trait evenness, and FDis index of trait divergence, calculated using three fluorescence channels, cell size from CLSM, as well as C assimilation, N assimilation, and S cell content) and genetic diversity metrics (Chao1 and Simpson index using 16S and 18S sequencing data) on cell density. Only metrics included in the list of models used for model averaging are shown here (predictors with a significant effect are in bold). Trait diversity (N=17) and genetic diversity (N=24) were modeled separately (see Methods). Values are standardized model-averaged regression coefficients shown as black circles with their 95% confidence intervals depicted as error bars (details about single models can be found in Extended Data Table S1). Cell density (quantified by scanning flow cytometry) is relative to the maximum achieved in a given layer.

### Single-cell traits drive niche and fitness differences in distinct lake layers

We analyzed our extensive single-cell dataset to investigate whether the highly competitive environment in deep layers of the lake drives increasing niche partitioning in pigment-related traits and decreasing fitness differences in metabolic traits. The data showed that the evenness of pigment-related traits was significantly higher in the chemocline (under low light intensity combined with high cell density) as compared to the upper layer (under highest light availability) (Fig. 4, Table 1A). The lower layer showed intermediate evenness of pigment-related traits, but differences with the upper layer and chemocline were not significant (Table 1A). In contrast, the evenness of metabolic traits did not show any significant difference between the three layers of the lake (Fig. 4 and Table 1B). This suggests that the evenness in traits related to light acquisition is positively related to increased niche partitioning under light limitation.

**Fig. 4:**
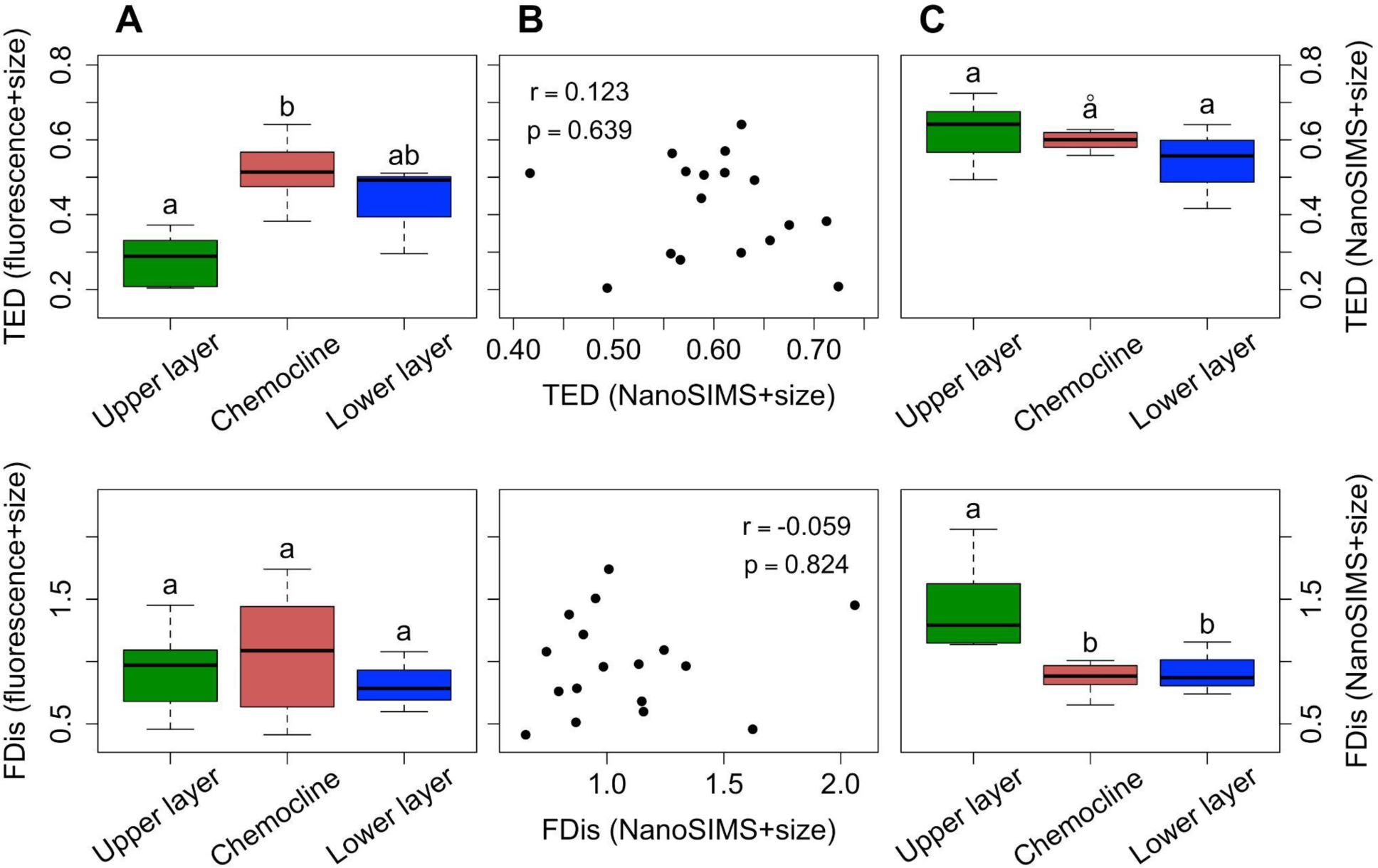
Niche and fitness differences are linked to different sets of traits. One-way ANOVA analyses and post hoc Tukey tests (see Table 1; significance level at p<0.05, N=17) show that the evenness of fluorescence traits (TED) is higher in the chemocline than in the upper layer, whereas trait divergence (FDis) does not show any significant difference (column **A**). Conversely, the evenness of metabolic traits measured with NanoSIMS does not change significantly between layers, but trait divergence is highest in the upper layer (column **C**). Fluorescence and metabolic trait distributions show weak and non-significant Spearman correlations (column **B**; function *cor.test*, N=17), further supporting the idea that fluorescence and metabolic traits reflect distinct mechanisms. Each boxplot reports minimum and maximum (whiskers), first and third quartile (lower and upper side, respectively), as well as median (horizontal black line). We note that, for the calculation of TED and FDis, cell size (from CLSM) was included along with three fluorescence channels from CLSM on one side (see Fig. 2A), as well as C assimilation, N assimilation and S cell content on the other side (see Fig. 2B).

**Table 1:** The results of the one-way ANOVA analyses and post hoc Tukey tests for the data shown in Fig. 4. All p-values are adjusted for multiple comparisons.

|  | **A) TED (fluorescence+size)** |  | **B) TED (NanoSIMS+size)** |  |
| --- | --- | --- | --- | --- |
|  | difference | p-value | difference | p-value |
| chemocline vs. upper layer | 0.235 [0.118; 0.351] | <0.001 | -0.015 [-0.119; 0.089] | 0.924 |
| lower layer vs. upper layer | 0.151 [-0.001; 0.303] | 0.052 | -0.086 [-0.222; 0.050] | 0.258 |
| lower layer vs. chemocline | -0.084 [-0.230; 0.062] | 0.319 | -0.071 [-0.201; 0.059] | 0.356 |

|  | **C) FDis (fluorescence+size)** |  | **D) FDis (NanoSIMS+size)** |  |
| --- | --- | --- | --- | --- |
|  | difference | p-value | difference | p-value |
| chemocline vs. upper layer | 0.123 [-0.451; 0.697] | 0.842 | -0.551 [-0.894; -0.207] | 0.002 |
| lower layer vs. upper layer | -0.117 [-0.868; 0.635] | 0.913 | -0.502 [-0.952; -0.052] | 0.028 |
| lower layer vs. chemocline | -0.240 [-0.960; 0.480] | 0.666 | 0.049 [-0.382; 0.479] | 0.953 |

The analysis further revealed that metabolic traits exhibit a significantly lower multidimensional metabolic trait divergence in the chemocline and lower layer than in the upper layer (Fig. 4 and Table 1D). In contrast, no significant difference in the divergence of pigment-related traits was observed between the three layers of the lake (Table 1C). This suggests that decreased divergence in metabolic traits related to resource acquisition is linked to decreased fitness differences. The latter occurred when competition for limiting resources (i.e., nutrients) increased. Taken together, our data suggest that niche partitioning and fitness effects act simultaneously between single-cell phenotypes in microbial communities along the depth gradient of Lake Cadagno and are linked to different sets of traits.

### Niche and fitness differences are negatively related because of resource limitation

After we confirmed that individual-level niche and fitness differences are evident in our dataset, we analyzed how they are linked to each other across the main environmental gradient of the lake. To this end, we formulated the hypothesis that single-cell niche and fitness differences are negatively related across depth. More specifically, our analysis showed that the cell-to-cell coefficient of variation in fitness proxies was negatively related to the evenness of traits related to pigments. This is arguably not unexpected because, with increasing depth, both increased niche partitioning of the light spectrum and decreased fitness differences due to more similar metabolic traits should occur. The results of the linear model (estimate= −2.378, SE=0.752, t=-3.164, df=15, R^2^=0.400; p=0.006) suggested that the individual-level niche partitioning of the light spectrum in deeper layers was strongly associated with more similar fitness as estimated by S content. Accordingly, cells in the upper layer were characterized by a high coefficient of variation in S content and uneven distribution of pigment-related traits (Fig. 5A, green dots), whereas cells in the chemocline and lower layer showed the expected combination of high evenness of pigment-related traits and low coefficient of variation in S content (Fig. 5A, red and blue dots). We also investigated the relationship between the evenness of pigment-related traits and the coefficient of variation in the single-cell assimilation rates of two other elements (C as CO_2_ and N as NH_4_^+^; Fig. 5B and C, respectively). We did not find any significant relationship between trait evenness of pigments and C assimilation rate (estimate=0.221, SE=1.537, t=0.144, df=15, R^2^=0.001; p=0.888), which suggests that CO_2_ assimilation is not a good proxy for fitness in this system. In contrast, the coefficient of variation of NH_4_^+^ assimilation rates showed a significant and positive relationship with the evenness of pigment-related traits (estimate=1.332, SE=0.584, t=2.280, df=15, R^2^=0.257; p=0.038). Cells in the upper layer (Fig. 5C, green dots) were generally more similar to each other in terms of NH_4_^+^ assimilation rates than cells in the chemocline and lower layer (Fig. 5C, red and blue dots).

**Fig. 5:**
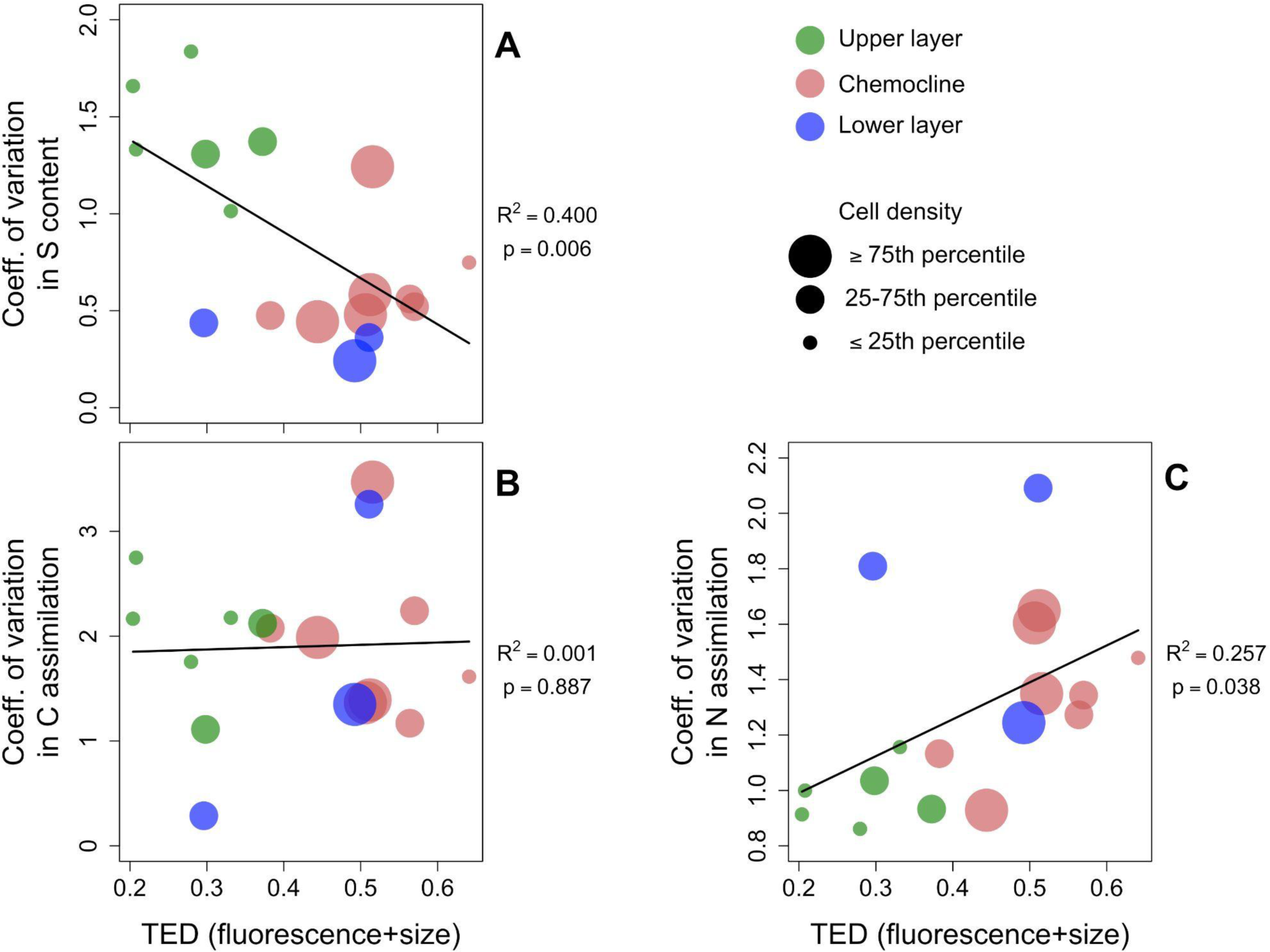
Relationship between niche and fitness differences across different layers of Lake Cadagno. Niche partitioning of the light spectrum was quantified using the TED index of trait evenness calculated with fluorescence data (three channels) plus cell size from CLSM data (see Fig. 2A & C). Fitness differences were quantified from NanoSIMS data (see Fig. 2B) using the coefficient of variation in S content (panel **A**), C assimilation (panel **B**) and N assimilation rates (panel **C**) among individual cells, whereby fitness differences decrease with decreasing coefficient of variation. Each circle represents a sample taken at a specific water depth and day of sampling. The color of the circles represents the association of the sample to a specific lake layer as defined in Fig. 1B. The size of the circle represents the association of each sample to a category of the magnitude of relative cell density in this layer as a proxy for ecosystem functioning. Black regression lines, R^2^, and p-values were obtained by fitting simple linear regressions (N=17).

## Discussion

In this study, trait diversity metrics integrating individual-level morphological, physiological and metabolic traits correlate significantly with ecosystem functioning, measured as cell densities, in contrast to taxonomic diversity metrics based on rRNA gene sequencing. This is consistent with previous findings: for example, multidimensional indices of functional diversity based on individual-based traits have been shown to outperform taxonomic diversity metrics in explaining phytoplankton biomass in natural communities^34^. Furthermore, the appropriate definition of microbial strategies (trait combinations) can help explain the contribution of microbial processes to carbon fluxes in soil ecosystems^48^. The lack of correlation between ecosystem functioning and taxonomic diversity, however, contradicts some previous findings^21^. One potential explanation for this is the importance of resource limitation in Lake Cadagno as compared to previously investigated systems. Previous studies have shown a positive relationship between resource limitation and intraspecific phenotypic heterogeneity in microbial populations^38,41,49^. Phenotypic heterogeneity can result from physiological acclimation and stochastic gene expression, and these physiological responses are not captured by taxonomic analysis. It is important to note that, in this case, physiology might have a key role in predicting ecosystem functioning, more than community composition or richness from taxonomic analysis. An alternative explanation could be that rRNA-based diversity has much greater interspecies redundancy than single-cell functional diversity for the traits we measured^50^ or that the measured traits show only a shallow phylogenetic conservation^51^. Indeed, to detect differences in trait diversity indices, it was necessary to first determine the minimum number of single cells per sample required for a robust analysis (Extended Data Fig. 2). It is noteworthy that 250 cells are sufficient for this purpose despite the presence of a high taxonomic diversity. This suggests that many different species and strains rely on similar functions and metabolic pathways^52^. Another explanation could be the extremely high accuracy with which we can measure differences in traits; for example, the consideration of the relative absorbance of multiple light wavelengths on a continuous scale allows us to distinguish individual cells based on slight differences in pigment profiles.

It has been recognized that trait variation among individuals must be considered to understand species coexistence^53,54^. These differences among individuals, attributable for example to phenotypic plasticity or phenotypic heterogeneity, affect both niche and fitness differences and therefore coexistence^30,55^. This may explain, for example, why the average cell or colony size of freshwater cyanobacterial species, while affecting both their niche and fitness differences, does not predict their coexistence even in simplified experimental communities^56^. This is consistent with our hypothesis that increasing niche partitioning (evenness in pigment-related traits) and decreasing fitness differences (convergence in metabolic traits) are better defined at a single-cell level when seeking a mechanistic understanding of microbial diversity. Niche and fitness differences are not necessarily independent but may be directly correlated, especially if they are both involved in metabolic activity and more specifically in photosynthesis. This interdependence should become apparent when using more specific and less integrative proxies for fitness. In this study, we find the cell content of S to be a good proxy for fitness, as sulfur globules ensure the availability of electron donors despite the extremely variable environmental conditions in the chemocline^57^. Indeed, sulfur bacteria can accumulate sulfur globules through sulfide uptake, and further use them as electron donors under conditions of sulfide limitation^58,59^. An even partitioning of the light spectrum among cells could also be directly responsible for the more similar values of cellular S (Fig. 5A). Indeed, this form of niche partitioning allows for a more uniform access to photons by decreasing competition for light, and the light-dependent accumulation of sulfur globules^58,59^. In contrast, N and C assimilation rates might not be good proxies for fitness, as a species dominating the uptake of both elements (i.e. *Chromatium okenii*) is present at very low densities in the chemocline of Lake Cadagno, whereas the most abundant species (i.e., *Chlorobium clathratiforme*) contributes much less to N and C cycles^57^. This may explain why we did not find any relationship between the evenness of pigment-related traits, our measure of niche partitioning, and the variability of C assimilation rates (Fig. 5B). On the contrary, N assimilation rates became increasingly variable among cells with increasing evenness of pigment-related traits (Fig. 5C). This suggests that sulfide (electron donor) limitation in the chemocline and lower layer drives heterogeneity in N assimilation, as previously shown experimentally for the fixation of N_2_ in a strain of the green sulfur bacterium *Chlorobium phaeobacteroides* isolated from Lake Cadagno^60^.

Our results support the idea that niche and fitness differences, even at the individual level, can be associated with different sets of traits, which can be measured simultaneously on the same single cells with the methods used in this study. In the chemocline and lower layer, cells show a higher evenness of pigment-related traits (i.e., niche partitioning of the light spectrum under light limitation, as observed in experimental phytoplankton populations^38^) and at the same time a lower variation in cell S content (i.e., more similar fitness). In other words, the environment of deep water layers, which we assume to be highly competitive because of high cell densities along with light and nutrient limitation, requires both high niche partitioning and low fitness differences for organisms to coexist. It should be noted that the strong correlation between these two mechanisms of coexistence (Fig. 5) does not imply a causal relationship. The fact that the cells increasingly partition the light spectrum might not necessarily cause the very similar cell S content but could otherwise indicate that S is a metabolic intermediate of an essential metabolic pathway in the chemocline and lower layer as opposed to the upper layer. Low S content variation in deep layers of the lake might reflect the role of S as an electron donor in anoxygenic photosynthesis. In order to survive there, cells need both enough photons to sustain photosynthesis (leading to high evenness in pigment-related traits) and enough S to channel their metabolism through this element (leading to low variation in cell S content).

Our findings have important implications for microbial ecology, biogeochemistry and community ecology. By measuring focal functional traits at the single-cell level, we connect the three above-mentioned fields in a novel conceptual framework that uses niche and fitness differences among individual cells to explain emerging properties at the community and ecosystem level, such as cell densities and standing biomass that sustains natural food webs. Fluorescence and metabolic traits represent fundamentally different axes of variation (see lack of correlation in Fig. 4), which only in combination can explain the occurrence of the corresponding phenotypes in a highly competitive environment. Our results further imply that, despite the high taxonomic complexity of natural microbial communities, single-cell trait-based approaches can explain community-level functions such as the nutrient cycling by reducing this complexity to a few relevant traits^24^, i.e., morphological, physiological and metabolic. Such strong links between traits and ecosystem properties across different environments have also been shown to exist at the level of microbiomes, aiding the identification of functional groups or guilds in metagenomic datasets^61,62^, as well as ecological clusters based on representative genomes^63^. Our approach might benefit the understanding of ecological communities in general, including those composed of larger organisms, and their links to broad-scale ecosystem functions^64^. Our findings support the need for manipulative experiments and emerging theoretical concepts that explain species coexistence using the fundamental unit at which interactions and selection occur in ecosystems, namely the individual organisms. Such concepts should eventually encompass all levels at which eco-evolutionary processes linked to environmental change occur: i.e. intraspecific trait variation resulting from phenotypic heterogeneity^30^ and phenotypic plasticity^55,65^, as well as rapid evolutionary adaptations^16^.

## Methods

### Study site, design, and sampling

The study was conducted in August 2014 in the meromictic lake ‘Lake Cadagno’ located in the Swiss Alps (1923 m above sea level) in Ticino, Switzerland. The maximum depth of the lake is 21 m. The water of the lake is infiltrated through gypsum-rich dolomite rock. The infiltrated water therefore transports salts, including sulfate, to the bottom water of the lake. This process establishes an anaerobic, euxinic monimolimnium with high salinity, and an aerobic, low-salinity mixolimnion separated by a permanent chemocline at ca. 10-14 m depth. The chemocline is characterized by sharp gradients and a strong turbidity maximum (ca. 1 m vertical thickness) dominated by populations of purple and green sulfur bacteria, which grow by anoxygenic photosynthesis with sulfide^57^. The exact depth of the turbidity maximum can fluctuate (between days or even within hours) as internal waves build up depending on the wind conditions.

Fieldwork was conducted for 3 consecutive days in August 2014. The days had varying sunlight exposure, introducing differences in photosynthetic activity. Each day, a multiparametric profile (temperature, conductivity, pH, salinity, dissolved oxygen, redox potential, and turbidity) was measured in the water column using a YSI 6000 profiler (Yellow Spring, Inc., Yellow Spring, OH). Water was retrieved through gas-tight tubing (Viton, d=6 mm) mounted to the multiparametric probe with a vacuum-driven pump (0.3 bar, 200 mL min^-1^) during the profile measurements. Samples were retrieved throughout the water column in 1 m steps from 1 to 10 m depth and in 0.5 m steps from 10.5 to 15 m depth. An overview on all samples and the associated analysis conducted with them is shown in Supplementary Dataset 1. Samples for stable isotope incubations were filled anoxically into N_2_-rinsed, septum-sealed, 250 mL serum bottles without head-space. Samples for chemical and flow cytometric analyses were sampled into 250 mL serum bottles that were cleaned by consecutive rinsing with 0.1% HCl and double-deionized water. Samples for DNA extraction and amplicon sequencing were sampled into 1 L bottles that were cleaned by consecutive rinsing with 2.5% NaClO, 0.1% HCl and 0.2 µm-filtered double-deionized water. Moreover, we measured a profile for photosynthetically active radiation as the percentage of transmission (PAR; Two LI-193SA spherical quantum sensors from LI-COR Ltd, Lincoln, NE) in the afternoon of each day after samples had been retrieved and stable isotope incubations had been placed in the lake.

### Chemical analysis

Samples for chemical analyses were filtered onsite through 0.47 µm nitrocellulose filters (Sartorius), stored cold and dark in autoclaved, crimp-sealed serum bottles without headspace, and analyzed within 10 days after sampling. Samples for ammonium were treated differently; these samples were filtered through 0.2 µm polycarbonate filters (GTTP, Millipore) and stored in 50 mL polypropylene tubes (consecutively rinsed with 0.1% HCl and 0.2 µm-filtered double-deionized water). Sulfate, chloride and nitrate were analyzed by ion chromatography (Metrohm 930 Compact IC Flex with chem. suppression; Metrohm Metrosep A Supp 5 100/4 mm column; conductivity detector). Ammonium was measured with Berthelot’s reaction (DIN 38406-5:1983-10). Ortho-phosphate was measured after the formation of a phosphorous-molybdenum blue-complex (DIN EN ISO 6878, 2004). Dissolved inorganic carbon was measured with TOC-L CSH (Shimadzu, Japan). Sulfide was measured one day from 12 mL samples that were immediately transferred to screw-capped tubes containing 0.8 mL 4% zinc acetate solution. These samples were stored in the dark and analyzed colorimetrically using a commercial kit (Spectroquant Sulfide kit, Merck, Schaffhausen, Switzerland). Total particulate nitrogen and carbon were determined by filtration of 100 to 220 mL lake water on pre-combusted (400°C, 3 h) 0.7-μm-pore glass microfibre filters (GF/F, Whatman) and subsequent measurement with isotope ratio mass spectrometry (IRMS). Briefly, the glass microfibre filters for IRMS were immediately stored at −20°C after filtration. After the fieldwork, samples were dried for at least 3 h at 65°C and decalcified in an atmosphere generated by 37% hydrochloric acid. The filters were kept dry in a desiccator at room temperature until further processing. The filters were measured by N_2_ and CO_2_ released by flash combustion in excess oxygen in an automated Thermo Flash EA 1112 elemental analyzer coupled to an isotopic ratio mass spectrometer (Thermo Delta Plus XP, Thermo Fisher Scientific). Standards were prepared from caffeine and total particulate carbon and nitrogen were calculated from the summed peak areas of masses 44, 45, and 46 for carbon and masses 28, 29, and 30 for nitrogen in each sample.

### Definition of lake layers

In order to determine the contribution of environmental factors to the distribution of samples in our dataset, we performed a principal component analysis (PCA; Fig. 1B). Individual samples were grouped into three major layers based on their position relative to the chemocline: upper layer, chemocline and lower layer. Samples from above the chemocline were further divided into a surface layer (initial 6 m of Lake Cadagno) and a layer just above the chemocline. This further subdivision was based on the fact that many environmental factors such as conductivity, temperature and oxygen concentrations were stable in the surface layer and gradually changed thereafter (Fig. 1A). In order to maximize differences between layers, we excluded from further analyses the layer between surface and chemocline, which shows intermediate properties and is therefore less clearly defined (Fig. 1B). For the PCA the following environmental factors were considered: depth [m], light [% PAR above the water], temperature [°C], conductivity [S/m], salinity [practical salinity units, psu], oxygen [mg L^-1^], pH, oxidation-reduction potential [mV], turbidity [formazine turbidity unit, FTU], ammonium [µmol N/L], dissolved inorganic carbon (DIC) [mg/L], ortho-phosphate [µg/L], chloride [mg/L], sulfate [mg/L], particulate nitrogen [µmolN/L], particulate carbon [µmolC/L]. For a small subset of points, individual values were not collected due to device failure. For these points, we calculated the mean of the respective variables using the samples directly above and below the missing value.

### Flow cytometry

Using the scanning flow cytometer Cytobuoy (www.cytobuoy.com, Woerden,The Netherlands), we counted and characterized phytoplankton cells from water sampled at different depths during each sampling day. Samples were fixed after stable isotope incubation with a fixative solution containing 0.01% paraformaldehyde and 0.1% glutaraldehyde at pH=7 and were stored at 4°C for 14 weeks until flow cytometric measurements. Cytobuoy contains two laser beams (coherent solid-state sapphire, wavelengths 488 and 635 nm) and three different detectors to measure the fluorescence emitted by photosynthetic pigments (ranges: 668–734 nm, 601–668 nm, 536–601 nm). Measurements were triggered by the sideward scattering signal. More details on the instrument can be found elsewhere^34^.

Cytometry data were successively cleaned by separating living phytoplankton cells from organic debris, suspended solids, and non-photosynthetic (heterotrophic) bacteria. First, a clustering algorithm was applied to a random subset of particles using the library *flowPeaks*^66^, and junk clusters were identified visually (characterized by low fluorescence in all pigments). Second, a random forest classifying algorithm^67,68^ was trained using the library *randomForest*^66^ to identify junk particles in the whole datasets and remove them. In addition, only particles in the size range between 1 μm and 1 mm are reliably characterized by Cytobuoy and were therefore retained in the dataset. The number of living phytoplankton cells identified was divided by the volume processed from each sample to calculate the cell density of photosynthetic cells as a proxy for primary production (ecosystem functioning).

### Stable isotope incubations

Stable isotope incubations were performed with anoxically-sampled, freshly-retrieved lake water as described under ‘Study site, design, and sampling’. The samples in the serum bottles were spiked with 100 µL stock solutions of ^15^N-NH_4_Cl (25 mM, Sigma) and ^13^C-NaHCO_3_ (0.5 M, Sigma) to a final concentration of 10 µM and 200 µM, respectively. The final labeling percentage (^13^C = 10 to 30%; ^15^N = 30 to 100%) was calculated for each sample retrospectively based on total NH_4_ and DIC measured through chemical analysis in separate water samples. After the addition of the isotopically-labeled substances, serum bottles were attached to a rope that was intruding into the water column and that was mounted on a stably-anchored research platform. The bottles were attached at different positions on the rope in a way that they incubated at the depth from which they had been retrieved. The bottles incubated for an entire day/night/day cycle (approximately 22 h) starting from about noon each day. The serum bottles were retrieved from the lake at the end of the incubations and kept in the dark and on ice until further processing (2 to 3 hours). Samples from the incubations were fixed with a fixative solution for later analysis with correlative imaging and flow cytometry (30 to 50 mL).

### Correlative imaging for autofluorescence and isotopic enrichment

Single-cell correlative imaging was performed with confocal laser scanning microscopy (CLSM, Leica SP5) for autofluorescence and nanometer-scale secondary ion mass spectrometry (NanoSIMS 50L, CAMECA) for isotopic enrichment within the Laboratory for Biological Geochemistry at EPFL and University of Lausanne. The cells were placed on polycarbonate filters (0.2 µm GTTP, Millipore) by vacuum filtration. The filters had a diameter of 5 mm. The diameter of the filtration area was 3 mm. Etched marks were distributed across the filtration area with laser micro-dissection (PALM micro-dissection, Zeiss 200M equipped with a 355 nm pulsed UV laser) before filtration. The filtered volumes of fixed cells were adjusted between 0.2 to 2 mL to have an evenly distributed, dense single-cell layer. Filtration of cells was immediately followed by filtration of 200 μL double-deionized water to wash the filters. Air-dried filters were coated with a thin film of low melting agarose by dipping them into a 0.1% low melting agarose solution at 35°C, removing excess agarose with a tissue by carefully touching the filter edge with the tissue, and air-drying the filters. The filters were stored dry and in the dark at room temperature until CLSM and NanoSIMS measurements.

First, the filters were imaged with CLSM to detect single-cell autofluorescence signals. To this end, cells were initially stained with a general DNA stain to enable cell detection. The filters were embedded in a solution containing the general DNA stain Hoechst (10 μg mL^−1^) and the antifading reagents Citifluor AF1 (Citifluor Ltd., PA; containing phosphate-buffered saline and glycerol) at 3 parts and Vectashield (Vecta Laboratories, CA) at 1 part. The cells on the embedded filters were imaged with CLSM by measuring z-stacks that were adjusted to extend from the filter surface until the top of the biggest cell including large algae and diatoms. For each sample, we obtained mosaics with 4 tiles from about 3 marks. The acquired images had the following dimensions: z-stack step-size of 2 µm (open pinhole) and xy-dimensions between 140 to 145 µm with a resolution of 1024×1024 pixels, resulting in a pixel size of about 0.14 µm (similar to the pixel size of NanoSims images). Images were recorded with bidirectional scanning at 400 Hz with a signal depth of 16 bit and averaging between 3 frames. Fluorescence was recorded in 17 channels varying in excitation (ex.; λ in nm) and emission (em.; λ in nm) during 9 sequential scans. Channels were selected based on (i) lambda scans of emitted light from samples obtained from different depths, (ii) spectra (in nm) of known photopigments (chlorophyll-a, chl-a; phycocyanin, PC; phycoerythrin, PE; fucoxanthin, Fuxo; accessory and decaying pigments, ADP) and (iii) equal fluorescence emission as recorded by the flow cytometer Cytobuoy. Scan 1: ex.: 633 / em.: 660-688 (chl-a/PC) & 725-750; scan 2: ex.: 633 / em.: 668-734 (Cytobuoy 1; chl-a/PE); scan 3: ex.: 543 / em.: 660-700 (PE/Fuxo); scan 4: ex.: 543 / em.: 625-650 & 700-750; scan 5: ex.: 488 / em.: 601-668 (Cytobuoy 2; PC) & 536-601 (Cytobuoy 3; ADP); scan 6: ex.: 488 / em.: 670-734 (chl-a/Fuxo) & 735-775; scan 7: ex.: 488 / em.: 518-548 & 565-605; scan 8: ex.: 458 / em.: 525-600 & 675-725 (chl-a) & reflected light; scan 9: ex.: 405 / em.: 423-503 (Hoechst stain) & reflected light. Imaging was done from higher to lower excitation wavelengths to reduce photobleaching. Gains were kept constant between all images. After imaging, the filters were washed in 100% ethanol and double-deionized water.

Next, the filters were prepared for NanoSIMS analysis by sputter-coating with a conductive gold layer (ca. 20 nm). Individual images or image mosaics were measured in the same regions that were imaged with CLSM previously. The respective areas were located using the etched marks and the CCD camera built into the NanoSIMS 50L. The measured areas were pre-sputtered with a Cs^+^ primary ion beam of about 4 pA to remove surface contamination, implant Cs^+^ ions, and achieve an approximately stable secondary ion emission rate. A primary Cs^+^ ion beam with a beam current between 1 and 1.2 pA and a beam diameter of around 150 nm was rastered across the cells for analysis with a dwell time of 5 ms per pixel. Secondary ion images for ^12^C^12^C^−^, ^13^C^12^C^−^, ^12^C^14^N^−^, ^12^C^15^N^−^, and ^32^S^−^ were simultaneously recorded from analysis areas of 30 μm × 30 μm with a resolution of 256×256 pixels, resulting in a pixel size of 0.117 µm. Ten planes from each individual area were measured. Mass resolving power was around 10,000 (Cameca definition), enough to resolve all measured isotopes from potential interferences in the mass spectrum.

Images were analyzed with a custom-written Matlab code that integrated Schnitzcells^69^ with Look@NanoSIMS^70^. First, a maximum intensity projection (MIP) image was created from the z-stack of the individual Hoechst images (scan 9). These MIP images, in which all cells were stained, were aligned with the corresponding NanoSIMS image in Look@NanoSIMS. The alignment was applied to all fluorescence images. Next, the aligned MIP image of scan 9 was used for automated segmentation of cells using Schnitzcells. This segmentation was manually corrected with an overlay image of Hoechst fluorescence and the ^12^C^14^N^−^ (biomass) NanoSIMS image for 268 to 662 cells per analyzed image. The segmentation was applied to all fluorescence images and the fluorescence intensities and cell dimensions for all channels were compiled. Finally, the segmentation was imported into Look@NanoSIMS to compile the corresponding isotope counts, isotopic fractions, and cell dimensions for each segmented cell. C and N assimilation for each segmented cell were calculated from their isotopic enrichment ^15^N/(^14^N+^15^N) and ^13^C/(^12^C+^13^C). Overall, images from 17 different samples coming from different days and depths were analyzed corresponding to a total of 6440 cells.

### Calculation of trait diversity

To obtain single-cell trait data, we combined stable isotope incubations - by adding ^15^N-NH_4_+ and ^13^C-CO_2_ into samples from different water depths and days - with correlative fluorescence microscopy and NanoSIMS (see previous two sections, as well as Fig. 2A and B). For the calculation of trait diversity, we first selected seven traits in total to avoid collinearity (Spearman correlation <0.7) and thus assure trait complementarity. Four traits were extracted from CLSM images: the mean fluorescence intensity of 3 channels: (i) excitation 543 nm; emission 700-750 nm, (ii), excitation 458 nm; emission 525-600 nm, (iii) excitation 458 nm; emission 675-725 nm, and (iv) cell size. Three traits were derived from NanoSIMS data: C and N assimilation, as well as S cell content. Those traits were then transformed to the same unit (mean=0, SD=1) and used to calculate three complementary indices of trait diversity, representing the distribution of individual cells of a microbial community in a high-dimensional trait space (Fig. 2C): TOP (*‘Trait Onion Peeling’*)^71^, TED (*‘Trait Even Distribution’*)^71^ and FDis (*‘Functional Dispersion’*)^72^. TOP, as a measure of trait richness, is sensitive to expansions and contractions of the trait space covered by the individuals of a community. TED, as a measure of trait evenness, is designed to respond to competitive interactions by quantifying niche partitioning as relative regularity in distances between individuals in the trait space. FDis, as a measure of trait divergence, reflects the degree to which individuals are spread around the average phenotype of the community. Please note that the list of traits included in the calculation of trait diversity varied according to the aim (and underlying hypotheses) of each single analysis, always being a selection of the above-mentioned seven traits (see ‘Statistical analysis’ below for more details). To robustly perform such analyses for a large array of samples and make trait diversity completely independent of the number of cells, we first had to determine the number of single cells in each sample that were required to detect differences in trait diversity indices between upper layer and chemocline (sensitivity analysis). To do this, we selected the two samples (A3 and A13) with the highest number of measured cells (662 and 540, respectively), and bootstrapped 100 times 50 to 450 cells (in intervals of 50) to calculate trait diversity indices (including all above mentioned seven traits). We then fitted linear regressions to test differences in trait diversity between the two samples. The data show that differences for each trait diversity index can be resolved by measuring approximately 250 cells (Extended Data Fig. 2). More specifically, goodness of fit and p-values indicate that at least 200 cells are necessary to distinguish upper layer and chemocline communities with a reasonable degree of confidence, especially with respect to the TED index of trait evenness (Extended Data Fig. 2). As the analyzed 17 samples had at least 269 cells, we decided to bootstrap 250 cells from each of them to further increase the representativeness of our subsamples. For further analyses, we considered the average trait diversity values of the 999 bootstrapped communities.

### Community analysis and calculation of genetic diversity

Samples for DNA extraction and amplicon sequencing were immediately filtered on 0.2 µm polycarbonate filters with a diameter of 25 mm (Millipore). The filtration volume was adjusted to filter approximately 10^7^ to 10^8^ cells on each filter, corresponding to 0.2 to 0.4 L per filter. Filters were stored at −20°C until further processing. An empty polycarbonate filter was processed as a negative control (DNA extraction and sequencing).

DNA was extracted from the filters using hot phenol/chloroform extraction. Cell lysis was performed with glass beads (106 µm) in 450 µL SDS-lysis buffer (35 mM sodium acetate, 7 mM EDTA, 2.9% SDS, pH 8) mixed with 1 mL phenol:chloroform:isoamyl alcohol (25:24:1 at pH 8) at 70°C for 12 min and bead beating for 2 min (Tissue Lyser II, Qiagen). The mixture was centrifuged at 4°C for 3 min at maximum speed and the aqueous phase was transferred to a new reaction tube. An equal volume of phenol:chloroform:isoamyl alcohol (25:24:1) was added to this tube. This extraction and centrifugation process was repeated for two more rounds. The final round of extraction was performed by adding chloroform:isoamyl alcohol (24:1) and again collecting the aqueous phase (∼200 µL). DNA was then precipitated from the recovered aqueous phase by adding 0.1 volume of 3 M sodium acetate (pH 8), gently mixing, and adding 2 volumes of 100 % ethanol and 1 µL of glycogen (20 mg/mL). Samples were incubated at 20°C for at least 3 hours, centrifuged at 4°C for 60 min at maximum speed and the supernatant was removed. The pellet was then washed twice with 200 µL of 80% ethanol, centrifuged at 4°C for 5 min and the supernatant removed. The pellet was dried under vacuum for 7 min and dissolved in 25 µL nuclease-free water. The DNA concentration was determined fluorometrically (Qbit, Invitrogen) and the concentration was normalized between samples to a maximum of 20 ng/µL by the addition of appropriate volumes of nuclease-free water.

Sequencing library preparation and DNA amplicon sequencing were performed by MR DNA (Molecular Research LP, Shallowater, TX) using a modified bTEFAP® library preparation protocol^73^ and sequencing on an Illumina MiSeq platform with a minimum of 20,000 reads per sample and read lengths of 2×300 bp. The 16S sequencing library was prepared with Bact341F: 5’-CCTACGGGNGGCWGCAG-3’ and Bact785R:5’-GACTACHVGGGTATCTAATCC-3’ ^74^, and the 18S sequencing library was prepared with euk516f:GGAGGGCAAGTCTGGT and euk1055r:CGGCCATGCACCACC^75^. A mock bacterial community was sequenced as a positive control (BEI Resources, Genomic DNA from Microbial Mock Community B, staggered, high Concentration, v5.2H, for Whole Genome Shotgun Sequencing, HM-277D).

We had to use two slightly different methods to process the sequence data of the two genes. The detailed protocol files describing the application, version, and parameters of the data processing for both genes can be found in the Appendix. In the first step, the sequence data were filtered to remove PhiX residues and sequences of low complexity. For the 16S amplicon data, read pairs were merged and primer site sequences were removed. For the 18S amplicon data, only the primer site trimmed forward reads were used as the reads could not be merged. We used UNOISE3^76^ to obtain zOTUs (zero-radius OTUs), which are considered equivalent to amplicon sequence variants (ASVs). We used SINTAX^76^ in combination with the reference SILVA SSU (v138 - https://www.arb-silva.de/) for 16S and pr2 SSU (v4.14.0 - https://pr2-database.org/) for 18S to predict the taxonomic affiliation of the zOTUs. The sequence data is deposited at NCBI/ENA under the BioProject number PRJNA1014378.

Finally, we calculated genetic diversity indices (Chao1 and Simpson) using the library *phyloseq*^66^ from unclustered zOTUs data.

### Statistical analysis

To test our first hypothesis, that ecosystem functioning is better explained by trait diversity than taxonomic diversity, we used a model averaging approach^77^. To this end, we formulated a list of linear models including as predictors all possible combinations (=16) of three trait diversity metrics (TOP, TED and FDis, calculated using three pigment-related traits, cell size, C assimilation, N assimilation, and S cell content) and day of sampling. In addition, in a second list of linear models we included as predictors all possible combinations (=32) of four genetic diversity metrics (Chao1 and Simpson index using 16S and 18S sequencing data) and the day of sampling. For both lists of models, the response variable was cell density relative to the maximum achieved in a given layer. Models with a difference in the Akaike information criterion corrected for small sample sizes (delta AICc<2) were defined as the most parsimonious set of models^78^ and were consequently used for model averaging (separately for trait and taxonomic diversity). Model selection and averaging were performed using the *MuMIn* R-package^66^. To address our second hypothesis that the highly competitive environment in deep layers of the lake drives increasing niche partitioning in pigment-related traits, and decreasing fitness differences in metabolic traits, we performed one-way ANOVA analyses and post hoc Tukey tests using the *TukeyHSD* function of the R-package *stats*. This procedure takes multiple comparisons into account when calculating p-values^66^. This allowed us to assess differences in the evenness of pigment-related traits and the divergence of metabolic traits between the three layers. To test our third hypothesis that single-cell niche and fitness differences are negatively related across depth, we fitted three simple linear models (*lm* function) with the evenness of pigment-related traits as a predictor and the coefficient of variation in S content and C and N assimilation rates as response variables.

## Acknowledgements

We thank the personnel of Piora Centro Biologia Alpina, especially Mauro Tonolla, for making it possible to use the laboratory facilities and housing during field work at lake Cadagno. We thank Martin Ackermann, Jukka Jokela and Dave Johnson (all Eawag) for discussions and support during the initial stages of the project. We thank Ilona Szivak (Eawag) for acquiring CLSM images. We thank the AUA chemical analysis teaching laboratory at Eawag for chemical analysis of sulfate, chloride, nitrate, ortho-phosphate, and dissolved inorganic carbon. We thank Nicola Storelli for reviewing the manuscript. This study was supported by Eawag, by the Swiss National Science Foundation Project 31003A_144053, an Eawag academic transition grant to F.S., a Synthesis Grant of the ETH Zurich Center for Adaptation to a Changing Environment (ACE) to F.S, and a Marie-Curie-Intra-European fellowship for career development (FP7-MC-IEF;271929;Phenofix) provided by the European Union to F.S. The NanoSIMS instrument in the Laboratory for Biological Geochemistry was funded in part by the European Union through a European Research Council Advanced Grant 246749 (BIOCARB) to A.M.

## Credit Authorship

**Simone Fontana:** Conceptualization, Methodology, Software, Formal analysis, Investigation, Data Curation, Writing - Original Draft, Writing - Review & Editing, Visualization, Project administration

**Désirée A. Schmitz:** Formal analysis (correlative imaging data), Data Curation, Writing - Original Draft, Writing - Review & Editing, Visualization

**Michael Daniels:** Formal analysis (PCA analysis), Writing - Review & Editing, Visualization

**Francesco Danza:** Investigation (field work), Resources, Writing - Review & Editing

**Thomas Röösli:** Software (for correlative image analysis), Writing - Review & Editing

**Hannah Bruderer:** Methodology, Investigation, Writing - Review & Editing

**Jean-Claude Walser:** Formal analysis (sequencing data), Data Curation, Writing - Review & Editing

**Julien Dekaezemacker:** Investigation (IRMS measurements), Writing - Review & Editing **Stephane Escrig:** Investigation (NanoSIMS measurements), Writing - Review & Editing **Anders Meibom:** Resources (NanoSIMS), Writing - Review & Editing, Supervision **Francesco Pomati:** Conceptualization, Writing - Review & Editing, Supervision

**Frank Schreiber:** Conceptualization, Methodology, Formal analysis, Data Curation, Writing - Original Draft, Writing - Review & Editing, Visualization, Supervision, Project administration, Funding acquisition

## References

1 Darwin, C. On the origin of species. (John Murray, 1859).

2 Grinnell, J. The Origin and Distribution of the Chest-Nut-Backed Chickadee. The Auk 21, 364–382 (1904).

3 Gause, G. F. Experimental Studies on the Struggle for Existence: I. Mixed Population of Two Species of Yeast. Journal of Experimental Biology 9, 389–402 (1932).

4 Grinnell, J. The Niche-Relationships of the California Thrasher. The Auk 34, 427–433 (1917).

5 Elton, C. Animal Ecology. (Sidgwick and Jackson, 1927).

6 Hutchinson, G. E. Concluding remarks. Cold Spring Harbor Symposia on Quantitative Biology 22, 415–427 (1957).

7 Hutchinson, G. E. The paradox of the plankton. The American Naturalist XCV, 137–145 (1961).

8 Chesson, P. Mechanisms of maintenance of species diversity. Annual Review of Ecology and Systematics 31, 343–366 (2000).

9 Chesson, P. Updates on mechanisms of maintenance of species diversity. Journal of Ecology 106, 1773–1794, doi:10.1111/1365-2745.13035 (2018).

10 Barabás, G., D’Andrea, R. & Stump, S. M. Chesson’s coexistence theory. Ecological Monographs 88, 277–303 (2018).

11 Cadotte, M. W. Concurrent niche and neutral processes in the competition-colonization model of species coexistence. Proceedings of the Royal Society B: Biological Sciences 274, 2739–2744, doi:10.1098/rspb.2007.0925 (2007).

12 Letten, A. D., Hall, A. R. & Levine, J. M. Using ecological coexistence theory to understand antibiotic resistance and microbial competition. Nature Ecology and Evolution 5, 431–441, doi:10.1038/s41559-020-01385-w (2021).

13 Song, C., Barabás, G. & Saavedra, S. On the Consequences of the Interdependence of Stabilizing and Equalizing Mechanisms. The American Naturalist 194, 627–639, doi:10.5061/dryad.j27h930 (2019).

14 Cordero, O. X. & Polz, M. F. Explaining microbial genomic diversity in light of evolutionary ecology. Nature Reviews Microbiology 12, 263–273, doi:10.1038/nrmicro3218 (2014).

15 Krismer, J., Tamminen, M., Fontana, S., Zenobi, R. & Narwani, A. Single-cell mass spectrometry reveals the importance of genetic diversity and plasticity for phenotypic variation in nitrogen-limited Chlamydomonas. The ISME Journal 11, 988–998, doi:10.1038/ismej.2016.167 (2017).

16 Hart, S. P., Turcotte, M. M. & Levine, J. M. Effects of rapid evolution on species coexistence. Proceedings of the National Academy of Sciences of the United States of America 116, 2112–2117, doi:10.1073/pnas.1816298116 (2019).

17 Ward, B. A. & Collins, S. Rapid evolution allows coexistence of highly divergent lineages within the same niche. Ecology Letters 25, 1839–1853, doi:10.1111/ele.14061 (2022).

18 Reiss, J., Bridle, J. R., Montoya, J. M. & Woodward, G. Emerging horizons in biodiversity and ecosystem functioning research. Trends in Ecology and Evolution 24, 505–514, doi:10.1016/j.tree.2009.03.018 (2009).

19 Eisenhauer, N. et al. Biodiversity–ecosystem function experiments reveal the mechanisms underlying the consequences of biodiversity change in real world ecosystems. Journal of Vegetation Science 27, 1061–1070, doi:10.1111/jvs.12435 (2016).

20 Turnbull, L. A., Levine, J. M., Loreau, M. & Hector, A. Coexistence, niches and biodiversity effects on ecosystem functioning. Ecology Letters 16, 116–127, doi:10.1111/ele.12056 (2013).

21 Patsch, D., van Vliet, S., Marcantini, L. G. & Johnson, D. R. Generality of associations between biological richness and the rates of metabolic processes across microbial communities. Environmental Microbiology 20, 4356–4368, doi:10.1111/1462-2920.14352 (2018).

22 Godoy, O., Gómez-Aparicio, L., Matías, L., Pérez-Ramos, I. M. & Allan, E. An excess of niche differences maximizes ecosystem functioning. Nature Communications 11, 4180–4180, doi:10.1038/s41467-020-17960-5 (2020).

23 Violle, C. et al. Let the concept of trait be functional! Oikos 116, 882–892, doi:10.1111/j.2007.0030-1299.15559.x (2007).

24 Fontana, S., Rasmann, S., de Bello, F., Pomati, F. & Moretti, M. Reconciling trait based perspectives along a trait-integration continuum. Ecology 102, e03472–e03472, doi:10.1002/ecy.3472 (2021).

25 Johnson, D. R. & Pomati, F. A brief guide for the measurement and interpretation of microbial functional diversity. Environmental Microbiology 22, 3039–3048, doi:10.1111/1462-2920.15147 (2020).

26 Lajoie, G. & Kembel, S. W. Making the Most of Trait-Based Approaches for Microbial Ecology. Trends in Microbiology 27, 814–823, doi:10.1016/j.tim.2019.06.003 (2019).

27 Achtman, M. & Wagner, M. Microbial diversity and the genetic nature of microbial species. Nature Reviews Microbiology 6, 431–440, doi:10.1038/nrmicro1872 (2008).

28 Philippot, L. et al. The ecological coherence of high bacterial taxonomic ranks. Nature Reviews Microbiology 8, 523–529, doi:10.1038/nrmicro2367 (2010).

29 Chase, A. B. et al. Emergence of soil bacterial ecotypes along a climate gradient. Environmental Microbiology 20, 4112–4126, 10.1111/1462-2920.14405 (2018).

30 Ackermann, M. A functional perspective on phenotypic heterogeneity in microorganisms. Nature Reviews Microbiology 13, 497–508, doi:10.1038/nrmicro3491 (2015).

31 Larkin, A. A. & Martiny, A. C. Microdiversity shapes the traits, niche space, and biogeography of microbial taxa. Environmental Microbiology Reports 9, 55–70, 10.1111/1758-2229.12523 (2017).

32 Bolnick, D. I. et al. Why intraspecific trait variation matters in community ecology. Trends in ecology & evolution 26, 183–192, doi:10.1016/j.tree.2011.01.009 (2011).

33 Krause, S. et al. Trait-based approaches for understanding microbial biodiversity and ecosystem functioning. Frontiers in Microbiology 5, 1–10, doi:10.3389/fmicb.2014.00251 (2014).

34 Fontana, S., Thomas, M. K., Moldoveanu, M., Spaak, P. & Pomati, F. Individual-level trait diversity predicts phytoplankton community properties better than species richness or evenness. ISME Journal 12, doi:10.1038/ismej.2017.160 (2018).

35 Hoppe, P., Cohen, S. & Meibom, A. NanoSIMS: Technical Aspects and Applications in Cosmochemistry and Biological Geochemistry. Geostandards and Geoanalytical Research 37, 111–154, 10.1111/j.1751-908X.2013.00239.x (2013).

36 Margalef, R. Life-forms of phytoplankton as survival alternatives in an unstable environment. Oceanologica Acta 1, 493–509, doi:10.1007/BF00202661 (1978).

37 Laughlin, D. C. & Messier, J. Fitness of multidimensional phenotypes in dynamic adaptive landscapes. Trends in Ecology and Evolution 30, 487–496, doi:10.1016/j.tree.2015.06.003 (2015).

38 Fontana, S., Thomas, M. K., Reyes, M. & Pomati, F. Light limitation increases multidimensional trait evenness in phytoplankton populations. ISME Journal 13, 1159–1167, doi:10.1038/s41396-018-0320-9 (2019).

39 Lemmen, K. D., Butler, O. M., Koffel, T., Rudman, S. M. & Symons, C. C. Stoichiometric traits vary widely within species: A meta-analysis of common garden experiments. Frontiers in Ecology and Evolution 7, 339–339, doi:10.3389/fevo.2019.00339 (2019).

40 Zaoli, S. et al. Generalized size scaling of metabolic rates based on single-cell measurements with freshwater phytoplankton. Proceedings of the National Academy of Sciences of the United States of America 116, 17323–17329, doi:10.1073/pnas.1906762116 (2019).

41 Schreiber, F. et al. Phenotypic heterogeneity driven by nutrient limitation promotes growth in fluctuating environments. Nature Microbiology 1, 16055–16055, doi:10.1038/nmicrobiol.2016.55 (2016).

42 Delong, J. P. & Hanson, D. T. Metabolic rate links density to demography in Tetrahymena pyriformis. ISME Journal 3, 1396–1401, doi:10.1038/ismej.2009.81 (2009).

43 Pettersen, A. K., White, C. R. & Marshall, D. J. Metabolic rate covaries with fitness and the pace of the life history in the field. Proceedings of the Royal Society B: Biological Sciences 283, 20160323–20160323, doi:10.1098/rspb.2016.0323 (2016).

44 Saini, J. S. et al. Bacterial, Phytoplankton, and Viral Distributions and Their Biogeochemical Contexts in Meromictic Lake Cadagno Offer Insights into the Proterozoic Ocean Microbial Loop. mBio 13, 1–20, doi:10.1128/mbio.00052-22 (2022).

45 Fox, J. W. & Vasseur, D. A. Character convergence under competition for nutritionally essential resources. American Naturalist 172, 667–680, doi:10.1086/591689 (2008).

46 Tonolla, M. et al. in Ecology of Meromictic Lakes (eds Ramesh D. Gulati, Egor S. Zadereev, & Andrei G. Degermendzhi) 155–186 (Springer International Publishing, 2017).

47 Tonolla, M., Peduzzi, S., Hahn, D. & Peduzzi, R. Spatio-temporal distribution of phototrophic sulfur bacteria in the chemocline of meromictic Lake Cadagno (Switzerland). FEMS Microbiology Ecology 43, 89–98, doi:10.1111/j.1574-6941.2003.tb01048.x (2003).

48 Malik, A. A. et al. Defining trait-based microbial strategies with consequences for soil carbon cycling under climate change. ISME Journal 14, 1–9, doi:10.1038/s41396-019-0510-0 (2020).

49 Schreiber, F. & Ackermann, M. Environmental drivers of metabolic heterogeneity in clonal microbial populations. Curr Opin Biotechnol 62, 202–211, doi:10.1016/j.copbio.2019.11.018 (2020).

50 Louca, S. et al. Function and functional redundancy in microbial systems. Nature Ecology and Evolution 2, 936–943, doi:10.1038/s41559-018-0519-1 (2018).

51 Martiny, J. B. H., Jones, S. E., Lennon, J. T. & Martiny, A. C. Microbiomes in light of traits: A phylogenetic perspective. Science 350, doi:10.1126/science.aac9323 (2015).

52 Johnson, D. R. et al. Association of biodiversity with the rates of micropollutant biotransformations among full-scale wastewater treatment plant communities. Applied and Environmental Microbiology 81, 666–675, doi:10.1128/AEM.03286-14 (2015).

53 Hart, S. P., Schreiber, S. J. & Levine, J. M. How variation between individuals affects species coexistence. Ecology Letters 19, 825–838, doi:10.1111/ele.12618 (2016).

54 Edwards, K. F. et al. Evolutionarily stable communities: a framework for understanding the role of trait evolution in the maintenance of diversity. Ecology Letters 21, 1853–1868, 10.1111/ele.13142 (2018).

55 Turcotte, M. M. & Levine, J. M. Phenotypic Plasticity and Species Coexistence. Trends in Ecology & Evolution 31, 803–813, doi:10.1016/j.tree.2016.07.013 (2016).

56 Gallego, I., Venail, P. & Ibelings, B. W. Size differences predict niche and relative fitness differences between phytoplankton species but not their coexistence. ISME Journal 13, 1133–1143, doi:10.1038/s41396-018-0330-7 (2019).

57 Musat, N. et al. A single-cell view on the ecophysiology of anaerobic phototrophic bacteria. Proceedings of the National Academy of Sciences of the United States of America 105, 17861–17866, doi:10.1073/pnas.0809329105 (2008).

58 Danza, F., Storelli, N., Roman, S., Lüdin, S. & Tonolla, M. Dynamic cellular complexity of anoxygenic phototrophic sulfur bacteria in the chemocline of meromictic Lake Cadagno. PLoS ONE 12, e0189510–e0189510, doi:10.1371/journal.pone.0189510 (2017).

59 Di Nezio, F. et al. Anoxygenic photo-and chemo-synthesis of phototrophic sulfur bacteria from an alpine meromictic lake. FEMS Microbiology Ecology 97, fiab010–fiab010, doi:10.1093/femsec/fiab010 (2021).

60 Zimmermann, M. et al. Substrate and electron donor limitation induce phenotypic heterogeneity in different metabolic activities in a green sulphur bacterium. Environmental Microbiology Reports 10, 179–183, doi:10.1111/1758-2229.12616 (2018).

61 Shan, X. & Cordero, O. X. Identifying microbial guilds on the basis of ecological patterns. Nature Ecology & Evolution 7, 651–652, doi:10.1038/s41559-023-02030-y (2023).

62 Shan, X., Goyal, A., Gregor, R. & Cordero, O. X. Annotation-free discovery of functional groups in microbial communities. Nature Ecology & Evolution 7, 716–724, doi:10.1038/s41559-023-02021-z (2023).

63 Guerra, C. A. et al. Global projections of the soil microbiome in the Anthropocene. Global Ecology and Biogeography 30, 987–999, doi:10.1111/geb.13273 (2021).

64 Eisenhauer, N. et al. The heterogeneity-diversity-system performance nexus. National Science Review 10, nwad109–nwad109, doi:10.1093/nsr/nwad109 (2023).

65 Hess, C., Levine, J. M., Turcotte, M. M. & Hart, S. P. Phenotypic plasticity promotes species coexistence. Nature Ecology & Evolution 6, 1256–1261, doi:10.1038/s41559-022-01826-8 (2022).

66 Team, R. C. in R Foundation for Statistical Computing (R Foundation for Statistical Computing, Vienna, Austria, 2022).

67 Breiman, L. Random forests. Machine Learning 45, 5–32, doi:10.1023/A:1010933404324 (2001).

68 Liaw, A. & Wiener, M. Classification and Regression by randomForest. R News 2, 18–22, doi:10.1177/154405910408300516 (2002).

69 Young, J. W. et al. Measuring single-cell gene expression dynamics in bacteria using fluorescence time-lapse microscopy. Nature Protocols 7, 80–88, doi:10.1038/nprot.2011.432 (2011).

70 Polerecky, L. et al. Look@NanoSIMS-a tool for the analysis of nanoSIMS data in environmental microbiology. Environmental Microbiology 14, 1009–1023, doi:10.1111/j.1462-2920.2011.02681.x (2012).

71 Fontana, S., Petchey, O. L. & Pomati, F. Individual-level trait diversity concepts and indices to comprehensively describe community change in multidimensional trait space. Functional Ecology 30, 808–818-808–818, doi:10.1111/1365-2435.12551 (2016).

72 Laliberté, E. & Legendre, P. A distance-based framework for measuring functional diversity from multiple traits. Ecology 91, 299–305 (2010).

73 Dowd, S. E., Sun, Y., Wolcott, R. D., Domingo, A. & Carroll, J. A. Bacterial Tag– Encoded FLX Amplicon Pyrosequencing (bTEFAP) for Microbiome Studies: Bacterial Diversity in the Ileum of Newly Weaned Salmonella-Infected Pigs. Foodborne Pathogens and Disease 5, 459–472, doi:10.1089/fpd.2008.0107 (2008).

74 Klindworth, A. et al. Evaluation of general 16S ribosomal RNA gene PCR primers for classical and next-generation sequencing-based diversity studies. Nucleic Acids Research 41, e1–e1, doi:10.1093/nar/gks808 (2013).

75 Stern, R. et al. Composition and Patterns of Taxa Assemblages in the Western Channel Assessed by 18S Sequencing, Microscopy and Flow Cytometry. Journal of Marine Science and Engineering 11, 480–480, doi:10.3390/jmse11030480 (2023).

76 Edgar, R. C. SINTAX: a simple non-Bayesian taxonomy classifier for 16S and ITS sequences. bioRxiv, 074161–074161, doi:10.1101/074161 (2016).

77 Burnham, K. P. & Anderson, D. R. Model Selection and Multimodel Inference. (Springer, 2002).

78 Symonds, M. R. E. & Moussalli, A. A brief guide to model selection, multimodel inference and model averaging in behavioural ecology using Akaike’s information criterion. Behavioral Ecology and Sociobiology 65, 13–21, doi:10.1007/s00265-010-1037-6 (2011).

