## Supplementary information for "Single-cell trait diversity explains niche and fitness differences in aquatic microbial communities"

This file contains extended data figures and a table for the article

[c]Kanton St. Gallen, Amt für Natur, Jagd und Fischerei, Abteilung Natur und Landschaft, St. Gallen, Switzerland.

[d]ETH Zurich, Dept. of Environmental Systems Science, Institute for Biogeochemistry and Pollutant Dynamics, Zurich, Switzerland.

[e]Eawag, Swiss Federal Institute for Aquatic Science and Technology, Dept. of Environmental Microbiology, Dübendorf, Switzerland.

[f]University of Zurich, Department of Quantitative Biomedicine, Zurich, Switzerland.

[g]University of Applied Sciences and Arts of Southern Switzerland (SUPSI), Department of Environment, Constructions and Design, Institute of Microbiology, Via Flora Ruchat-Roncati 15, 6850 Mendrisio, Switzerland.

[h]ETH Zurich, Dept. of Environmental Systems Science, Genetic Diversity Centre (GDC), Zurich, Switzerland.

[i]Max Planck Institute for Marine Microbiology, Dept. of Biogeochemistry, Bremen, Germany.

[j]Laboratory for Biological Geochemistry, School of Architecture, Civil and Environmental Engineering (ENAC), École Polytechnique Fédérale de Lausanne (EPFL), Lausanne, Switzerland.

[k]Center for Advanced Surface Analysis, Institute of Earth Sciences, University of Lausanne, Lausanne, Switzerland.

[l]Federal Institute for Materials Research and Testing (BAM), Department of Materials and Environment, Division of Biodeterioration and Reference Organisms, Berlin, Germany.

The supplementary information comprises:

- 2 extended data figures: Figure S1 – S2
- 1 extended data table: Table S1

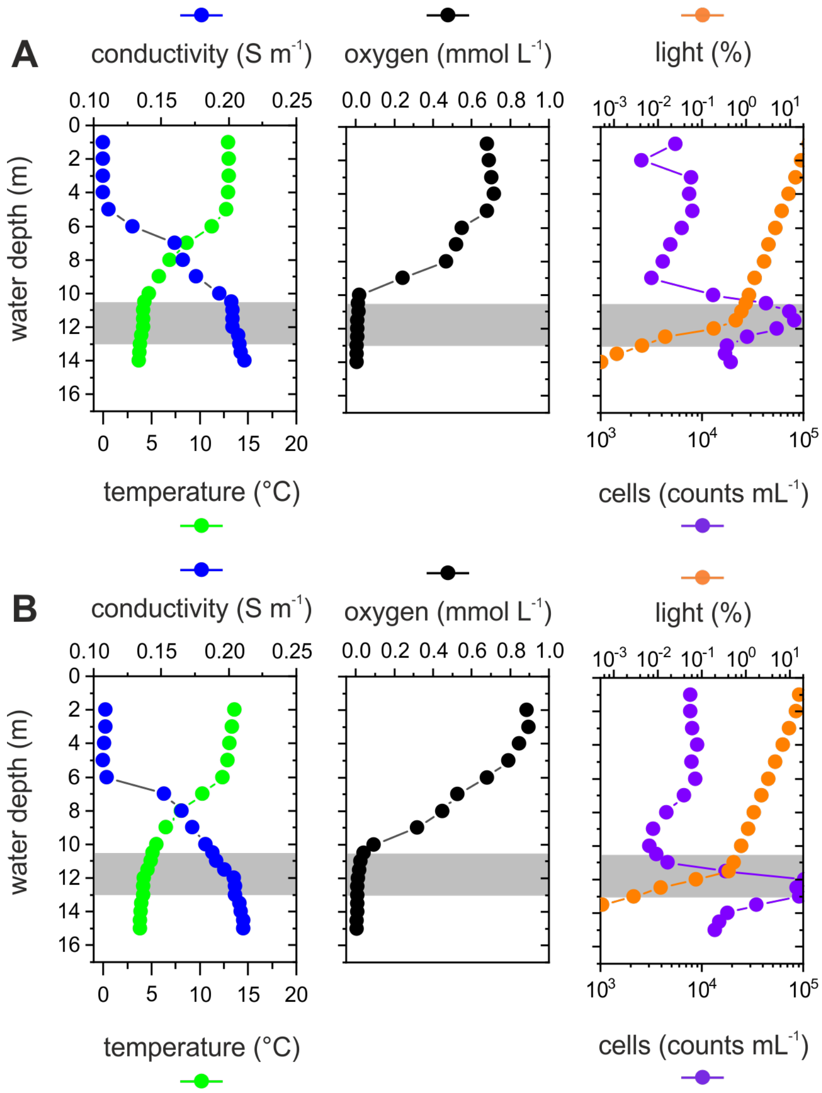


**Extended Data Fig. S1.** Vertical gradients of physical, chemical and biological parameters in Lake Cadagno separate the ecosystem into three distinct layers. Vertical profiles of conductivity (blue), temperature (green), oxygen (black), total sulfide (red), light (orange), and cell density (purple) measured on one day of the sampling campaign. Panel **(A)** shows profiles from the sampling day A (17/08/2014) and panel **(B)** shows profiles from sampling day B (18/08/2014). Data from sampling day C (19/08/2014) are shown in Fig. 1. The gray area represents the chemocline, where sulfide and oxygen are depleted, the cell density increases, and light gets severely limited. Light values are presented as % photosynthetic active radiation of the light above the water. Light values shown as half circles below the chemocline were below the detection limit of the sensor.

(A)

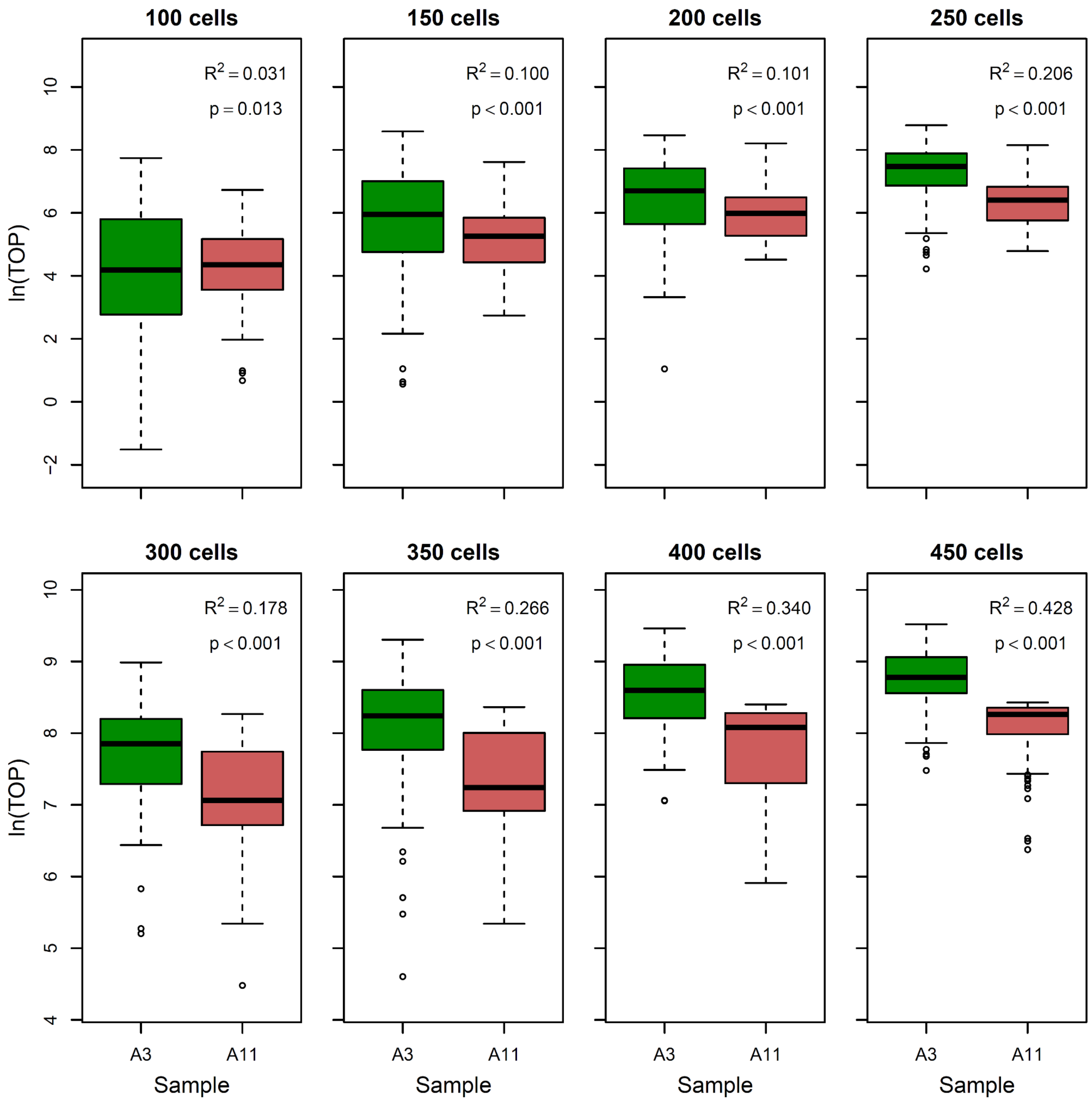
100 cells
150 cells
200 cells
250 cells
300 cells
350 cells
400 cells
450 cells
$R^2 = 0.031$
p = 0.013
$R^2 = 0.100$
p < 0.001
$R^2 = 0.101$
p < 0.001
$R^2 = 0.206$
p < 0.001
$R^2 = 0.178$
p < 0.001
$R^2 = 0.266$
p < 0.001
$R^2 = 0.340$
p < 0.001
$R^2 = 0.428$
p < 0.001
ln(TOP)
A3
A11
Sample

(B)

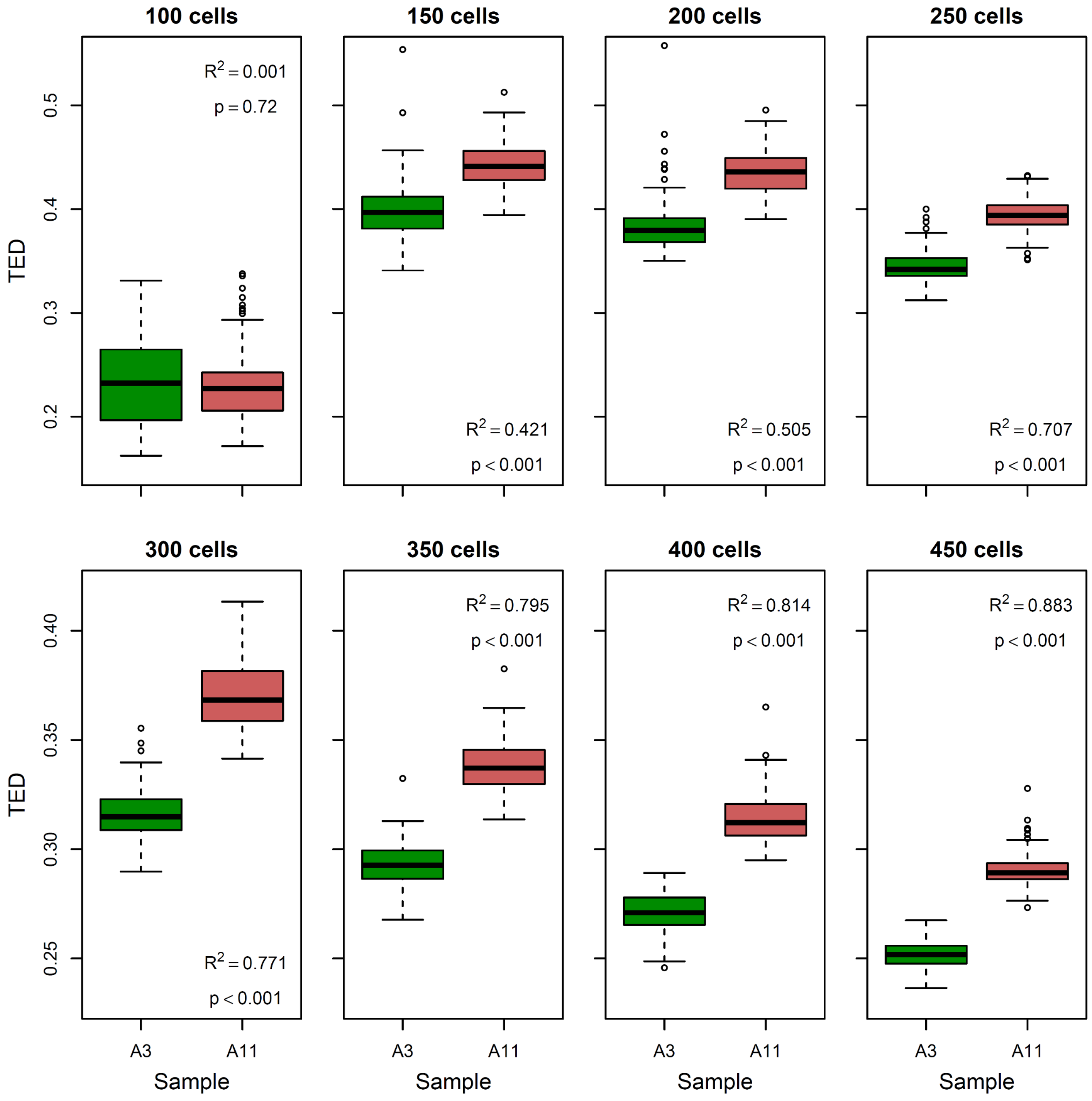
100 cells
150 cells
200 cells
250 cells
300 cells
350 cells
400 cells
450 cells
TED
TED
Sample
A3
A11
R2 = 0.001
p = 0.72
R2 = 0.421
p < 0.001
R2 = 0.505
p < 0.001
R2 = 0.707
p < 0.001
R2 = 0.771
p < 0.001
R2 = 0.795
p < 0.001
R2 = 0.814
p < 0.001
R2 = 0.883
p < 0.001
0.2
0.3
0.4
0.5
0.25
0.30
0.35
0.40

**(C)**

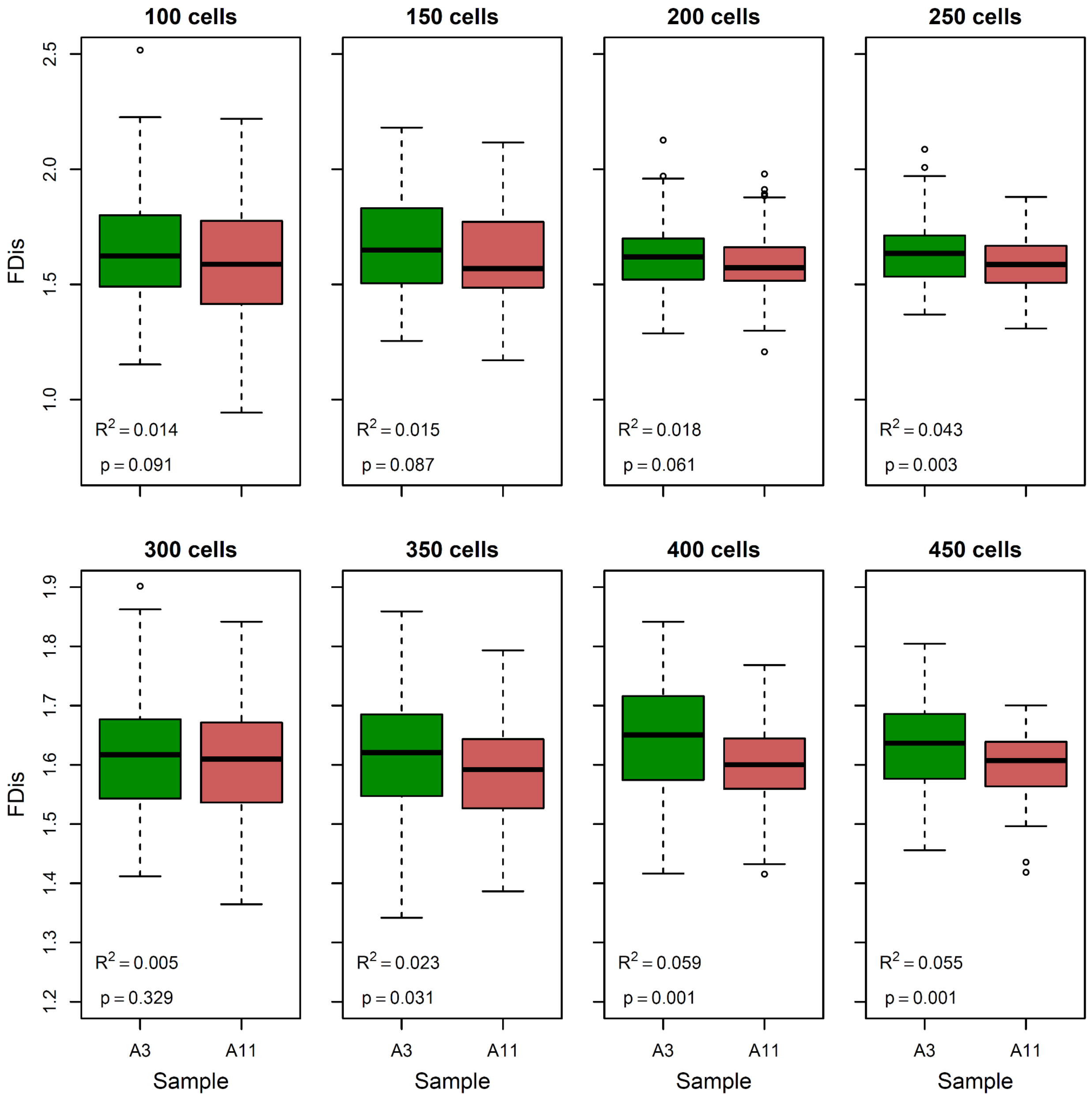


**Extended Data Fig. S2:** Sensitivity analysis to test how many cells must be sampled to detect differences in **(A)** trait richness (TOP), **(B)** trait evenness (TED), and **(C)** trait divergence (FDis) between upper layer (sample A3) and chemocline (sample A11). A number of cells ranging from 50 to 450 (in intervals of 50) was randomly sampled from A3 and A11 (100 times each), and linear regressions were fitted to assess differences between the two groups ($R^2$ and p-values are reported in the graphs). Note that TOP values were transformed with the natural logarithm only to ease the visualization but not in the underlying models.

**Extended Data Table S1:** List of all models with trait diversity and genetic diversity metrics (as well as Day) as predictors of cell density (relative to the maximum achieved in a given layer). The models are ranked according to AICc values, and gray shading indicates the subset of models used for model averaging (delta AICc < 2; see Fig. 3). Standardized regression estimates are indicated.

| Trait diversity | Intercept | TOP | TED | FDis | Day | df | logLik | AICc | delta | weight | $R^2$ |
|---|---|---|---|---|---|---|---|---|---|---|---|
| | 0.649 | | -0.217 | 0.149 | | 4 | 2.04 | 7.26 | 0.00 | 0.380 | 0.568 |
| | 0.642 | | -0.143 | 0.164 | + | 6 | 5.71 | 8.98 | 1.72 | 0.161 | 0.719 |
| | 0.649 | | -0.205 | | | 3 | -1.12 | 10.08 | 2.83 | 0.092 | 0.373 |
| | 0.649 | 0.113 | -0.217 | | | 4 | 0.56 | 10.21 | 2.96 | 0.087 | 0.485 |
| | 0.671 | | | 0.172 | + | 5 | 2.46 | 10.54 | 3.29 | 0.074 | 0.588 |
| | 0.649 | 0.014 | -0.217 | 0.139 | | 5 | 2.06 | 11.34 | 4.09 | 0.049 | 0.568 |
| | 0.618 | 0.147 | -0.140 | | + | 6 | 4.51 | 11.38 | 4.12 | 0.048 | 0.677 |
| | 0.647 | 0.158 | | | + | 5 | 1.73 | 11.99 | 4.73 | 0.036 | 0.552 |
| | 0.581 | | -0.154 | | + | 5 | 0.98 | 13.50 | 6.24 | 0.017 | 0.510 |
| | 0.609 | | | | + | 4 | -1.33 | 13.99 | 6.74 | 0.013 | 0.357 |
| | 0.642 | 0.063 | -0.140 | 0.120 | + | 7 | 6.22 | 14.01 | 6.76 | 0.013 | 0.735 |
| | 0.670 | 0.074 | | 0.121 | + | 6 | 2.93 | 14.54 | 7.28 | 0.010 | 0.611 |
| | 0.649 | | | | | 2 | -5.09 | 15.03 | 7.77 | 0.008 | 0.000 |
| | 0.649 | | | 0.133 | | 3 | -3.65 | 15.14 | 7.89 | 0.007 | 0.156 |
| | 0.649 | 0.092 | | | | 3 | -4.43 | 16.70 | 9.45 | 0.003 | 0.075 |
| | 0.649 | -0.007 | | 0.137 | | 4 | -3.65 | 18.63 | 11.37 | 0.001 | 0.156 |

| Genetic diversity | Intercept | Chao1 (18S) | Chao1 (16S) | Simpson (18S) | Simpson (16S) | Day | df | logLik | AICc | delta | weight | $R^2$ |
|---|---|---|---|---|---|---|---|---|---|---|---|---|
| | 0.698 | | | | | | 2 | -5.33 | 15.22 | 0.00 | 0.281 | 0.000 |
| | 0.698 | | | 0.073 | | | 3 | -4.63 | 16.46 | 1.23 | 0.152 | 0.057 |
| | 0.698 | | | | -0.040 | | 3 | -5.13 | 17.45 | 2.23 | 0.092 | 0.016 |
| | 0.698 | | -0.036 | | | | 3 | -5.17 | 17.53 | 2.31 | 0.089 | 0.013 |
| | 0.698 | 0.024 | | | | | 3 | -5.25 | 17.71 | 2.48 | 0.081 | 0.006 |
| | 0.698 | | -0.048 | 0.081 | | | 4 | -4.32 | 18.75 | 3.52 | 0.048 | 0.080 |
| | 0.698 | | | 0.071 | -0.035 | | 4 | -4.47 | 19.04 | 3.81 | 0.042 | 0.069 |
| | 0.698 | -0.029 | | 0.090 | | | 4 | -4.55 | 19.21 | 3.99 | 0.038 | 0.062 |
| | 0.715 | | | | | + | 4 | -4.85 | 19.81 | 4.58 | 0.028 | 0.039 |
| | 0.698 | 0.036 | | | -0.048 | | 4 | -4.97 | 20.05 | 4.82 | 0.025 | 0.029 |
| | 0.698 | | -0.031 | | -0.036 | | 4 | -5.00 | 20.11 | 4.88 | 0.024 | 0.027 |
| | 0.698 | 0.033 | -0.043 | | | | 4 | -5.03 | 20.17 | 4.94 | 0.024 | 0.024 |
| | 0.698 | | -0.044 | 0.078 | -0.029 | | 5 | -4.21 | 21.74 | 6.52 | 0.011 | 0.089 |
| | 0.698 | -0.021 | -0.046 | 0.093 | | | 5 | -4.28 | 21.90 | 6.67 | 0.010 | 0.083 |
| | 0.698 | -0.016 | | 0.081 | -0.030 | | 5 | -4.45 | 22.22 | 7.00 | 0.008 | 0.071 |
| | 0.708 | | | 0.058 | | + | 5 | -4.49 | 22.31 | 7.09 | 0.008 | 0.067 |
| | 0.732 | | | | -0.051 | + | 5 | -4.53 | 22.39 | 7.17 | 0.008 | 0.064 |
| | 0.709 | | -0.031 | | | + | 5 | -4.72 | 22.78 | 7.56 | 0.006 | 0.049 |
| | 0.698 | 0.044 | -0.039 | | -0.046 | | 5 | -4.78 | 22.89 | 7.66 | 0.006 | 0.045 |
| | 0.707 | 0.021 | | | | + | 5 | -4.80 | 22.93 | 7.71 | 0.006 | 0.043 |
| | 0.698 | -0.010 | -0.044 | 0.084 | -0.027 | | 6 | -4.20 | 25.34 | 10.11 | 0.002 | 0.090 |
| | 0.698 | | -0.045 | 0.068 | | + | 6 | -4.23 | 25.41 | 10.18 | 0.002 | 0.087 |
| | 0.724 | | | 0.051 | -0.044 | + | 6 | -4.25 | 25.43 | 10.21 | 0.002 | 0.086 |
| | 0.722 | 0.032 | | | -0.057 | + | 6 | -4.40 | 25.74 | 10.52 | 0.001 | 0.074 |
| | 0.726 | | -0.025 | | -0.047 | + | 6 | -4.45 | 25.84 | 10.62 | 0.001 | 0.070 |
| | 0.714 | -0.022 | | 0.073 | | + | 6 | -4.45 | 25.85 | 10.62 | 0.001 | 0.070 |
| | 0.696 | 0.030 | -0.039 | | | + | 6 | -4.61 | 26.17 | 10.95 | 0.001 | 0.058 |
| | 0.713 | | -0.038 | 0.061 | -0.037 | + | 7 | -4.07 | 29.13 | 13.91 | 0.000 | 0.100 |
| | 0.702 | -0.014 | -0.043 | 0.078 | | + | 7 | -4.22 | 29.44 | 14.21 | 0.000 | 0.088 |
| | 0.724 | -0.001 | | 0.051 | -0.044 | + | 7 | -4.25 | 29.49 | 14.27 | 0.000 | 0.086 |
| | 0.712 | 0.040 | -0.033 | | -0.054 | + | 7 | -4.26 | 29.52 | 14.30 | 0.000 | 0.085 |
| | 0.712 | 0.004 | -0.038 | 0.058 | -0.038 | + | 8 | -4.07 | 33.73 | 18.51 | 0.000 | 0.100 |
